# MAX inactivation deregulates the MYC network and induces neuroendocrine neoplasia in multiple tissues

**DOI:** 10.1101/2024.09.21.614255

**Authors:** Brian Freie, Ali H. Ibrahim, Patrick A. Carroll, Roderick T Bronson, Arnaud Augert, David MacPherson, Robert N. Eisenman

**Affiliations:** Basic Sciences Division, Fred Hutchinson Cancer Center, Seattle WA USA; Human Biology and Public Health Science Divisions, Fred Hutchinson Cancer Center, Seattle WA USA; Division of Immunology, Department of Microbiology and Immunobiology, Harvard Medical School, Boston, MA, USA; Department of Genome Sciences, University of Washington, Seattle WA USA; Present address: Yale Cancer Center, New Haven, CT 06520, USA; Department of Pathology, Yale School of Medicine, Yale University, New Haven, CT 06510, USA; Present address: Department of Internal Medicine, The University of Texas Health Science Center, Houston TX USA

## Abstract

The MYC transcription factor requires MAX for DNA binding and widespread activation of gene expression in both normal and neoplastic cells. Surprisingly, inactivating mutations in *MAX* are associated with a subset of neuroendocrine cancers including pheochromocytoma, pituitary adenoma and small cell lung cancer. Neither the extent nor the mechanisms of MAX tumor suppression are well understood. Deleting *Max* across multiple mouse neuroendocrine tissues, we find *Max* inactivation alone produces pituitary adenomas while *Max* loss cooperates with *Rb1*/*Trp53* loss to accelerate medullary thyroid C-cell and pituitary adenoma development. In the thyroid tumor cell lines, MAX loss triggers a striking shift in genomic occupancy by other members of the MYC network (MNT, MLX, MondoA) supporting metabolism, survival and proliferation of neoplastic neuroendocrine cells. Our work reveals MAX as a broad suppressor of neuroendocrine tumorigenesis through its ability to maintain a balance of genomic occupancies among the diverse transcription factors in the MYC network.

**Teaser:** *MAX* inactivation deregulates multiple transcription factors to induce neuroendocrine cancers

## Introduction

The basic-helix-loop-helix-zipper (bHLHZ) protein, MAX, is the obligate physiological heterodimeric binding partner for the three members of the MYC protein family as well as for other members of the proximal MYC bHLHZ network (MXD1-4, MNT and MGA) (*1–6*). Dimerization with MAX permits the widespread binding of MYC-MAX to E-Box DNA and activation of multiple transcriptional programs relevant to cell growth, metabolism, apoptosis and proliferation (*7–13*). Research from many labs has confirmed that MYC exists predominantly as a heterodimer with MAX *in vivo*, and that MYC-MAX dimerization is required for MYC function in normal as well as neoplastic cells harboring deregulated MYC (*14–17*). The essentiality of MAX is underscored by the early embryonic lethality associated with constitutive homozygous *Max* deletion in mice (*17, 18*). While a MAX-like protein known as MLX has been identified and possesses a highly related bHLHZ domain, it does not interact with either MAX or MYC but can heterodimerize with MNT and the large cytoplasmic bHLHZ proteins MondoA (MLXIP) and ChREBP (MLXIPL) (*19–21*). Through these interactions, MLX has been shown to influence metabolism and cell stress in a manner distinct from MAX (*22, 23*). Moreover, in contrast to *Max* loss-of-function, constitutive homozygous deletion of *Mlx* does not compromise normal murine development, postnatal viability or lifespan, although it does result in male-only infertility related to intrinsic activation of apoptosis during spermatogenesis (*24*).

One long-standing puzzle pertaining to MAX is that, despite the copious evidence that it is essential for MYC’s activity as a fundamental driver of oncogenesis, MAX loss of function occurs in a significant subset of neuroendocrine neoplasms, suggestive of MAX function as a tumor suppressor. Initially, the rat pheochromocytoma PC12 cell line was reported to lack MAX (*25*). Later, germline inactivating mutations in *Max* were discovered in hereditary pheochromocytomas (*26, 27*) and in neuroendocrine pituitary tumors (*28*) while sporadic *Max* mutations are found in small cell lung cancer (SCLC) (*29*), GIST and other neuroendocrine tumors (*30, 31*). Our studies, using mouse models, demonstrated that while homozygous *Max* deletion completely abrogates lymphomagenesis in *Eμ-Myc* mice, the same *Max* deletion in lung epithelium cooperates with *Rb1/Trp53* loss to promote SCLC (*17, 32*). In both models, MAX loss triggered disappearance of MYC most likely due to its increased degradation, consistent with other reports that MYC stability is in part controlled by its interaction with MAX (*33–35*). Furthermore, in cells harboring *Max* deletions, genomic binding by MAX, MYC and MNT was dramatically reduced (*32*). We also demonstrated that *Max* loss led to de-repression of metabolic one-carbon pathway genes and to an increase in serine biosynthesis, leading to survival in the absence of external serine (*32*).

While germline inactivating mutations in *MAX* occur in neuroendocrine cancers such as paraganglioma, pheochromocytoma (*26, 27*) and, with lower penetrance, pituitary tumorigenesis (*28, 36*), the actual role that MAX loss of function plays in these tumor types has not been studied and no mouse models are available. Many of the tumor types harboring *MAX* mutations/deletions are neuroendocrine, prompting us to delete *Max* across many neuroendocrine cell compartments in a mouse model and to assess impact on tumorigenesis and changes in gene expression and genomic occupancy by members of the MYC network that are critical for neoplastic initiation and progression.

## Results

We crossed our conditional *Max*-floxed allele into an Ascl1-CreERT2 strain (*17, 37*) in which an inducible form of CreERT2 was knocked into the endogenous *Ascl1* locus. Tamoxifen was delivered to adult mice, with injections over 5 consecutive days to drive *Max* deletion across different neuroendocrine compartments (Fig. 1A). When mice became moribund, we performed necropsy analyses. As shown in Fig 1B, *Ascl1CreERT2 Max^lox/lox^*animals exhibited median survival of 604 days post tamoxifen delivery. Histological analyses revealed pituitary adenomas in the *Max-*deleted mice (8 of 13 mice) but not in controls (Fig. 1C). Western blot analysis shows pituitary tumors from mice with an *Rb1/Trp53* deletion (wild-type *Max*) compared with the tumors arising from *Max* deletion alone. The latter exhibit loss of the full-length MAX protein, and the appearance of a faster migrating band consistent with the MAX N-terminal fragment resulting from the targeted deletion of the MAX helix2-zipper region required for dimerization with MYC (*17*) (Fig. 1D) (see Discussion). These data are the first to demonstrate that *Max* inactivation alone is sufficient to initiate tumorigenesis in mice.

**Figure 1:**
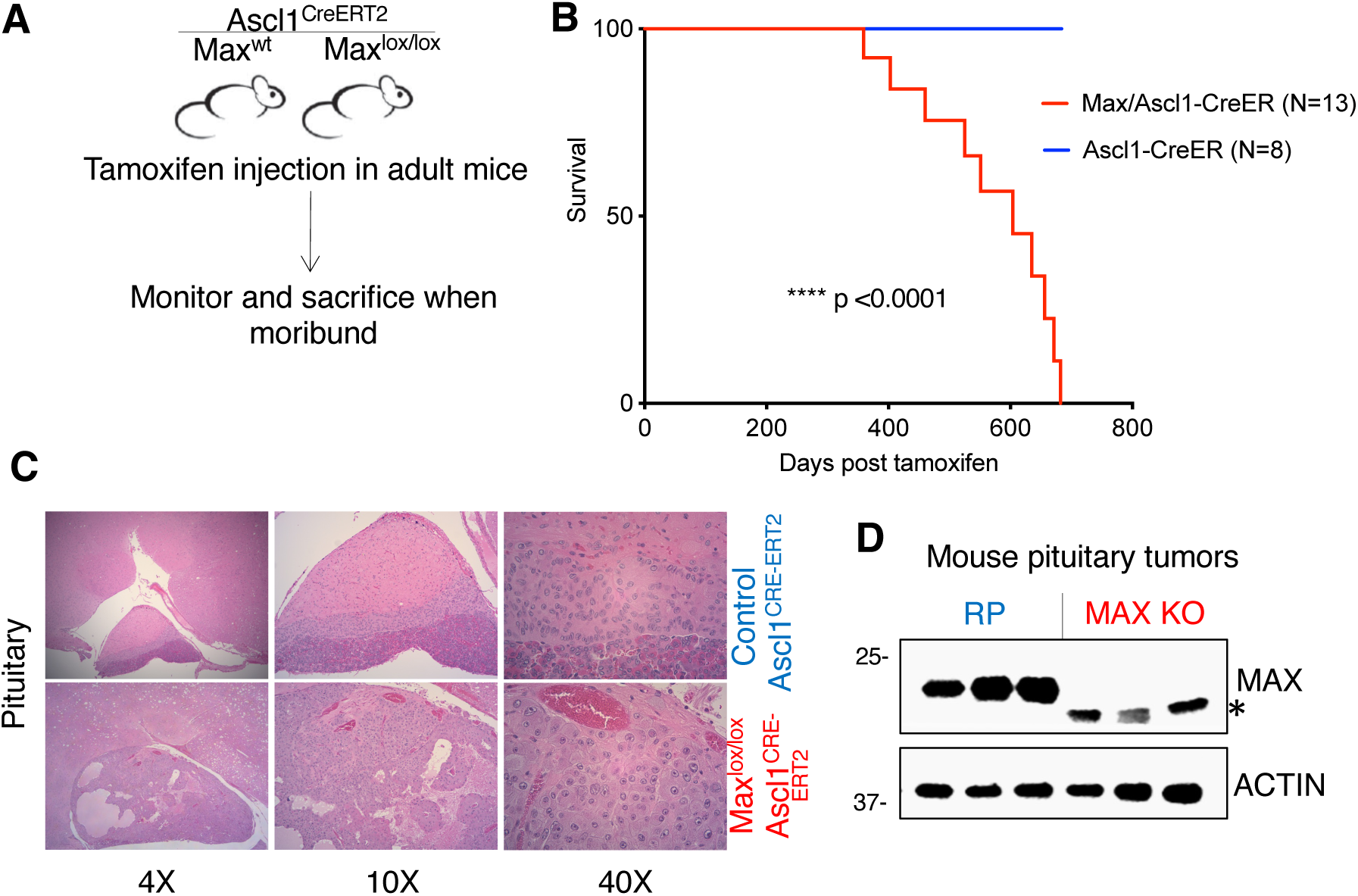
*Max* deletion results in pituitary adenomas that arise with long latency. **A)** Illustration of mouse models employed to study roles for *Max* in neuroendocrine tumorigenesis. **B)** Kaplan-Meier curve showing survival with *Max* inactivation in the Ascl1-Cre-ERT2 model following tamoxifen delivery with p-value from log-rank test shown. **C)** Histology showing pituitary lesions in control and *Max*-deleted mice. **D)** Western blot analyses of pituitary tumors arising with *Max* deletion. As a positive control, pituitary tumors from RP mice with *Rb1/Trp53* deletion driven by Ascl1-Cre-ERT2 are also shown. Asterisk indicates position of the N-terminal fragment of the truncated MAX protein in RPMax cells.

The incomplete penetrance and long latency of pituitary tumor formation seen in this model limited our ability to study key aspects of MAX function in neuroendocrine tumor suppression. Thus, we turned to a sensitized genetic background with *Rb1/Trp53* deletion (RP mice) that we previously used to study the impact of tumor suppressor gene deletion on neuroendocrine cancer types (*38*). We bred the *Ascl1CreERT2 Max^lox/lox^* mice into an *Rb1*/*Trp53* floxed background, again delivering tamoxifen to adult mice over 5 consecutive days (Fig. 2A). We performed both long-term tumor survival studies and we also examined phenotypes at a common time point, 10 weeks post tamoxifen delivery. At the 10-week time point, the RP mice exhibited hyperplastic lesions in the thyroid gland (Fig. 2B), consistent with *Rb1/Trp53* loss promoting thyroid tumorigenesis in mice (*39*). In contrast to the observed hyperplastic lesions that we observed in the RP model, every RPMax mouse developed bilateral thyroid C-cell adenomas. While *Rb1/Trp53* loss leads to neuroendocrine pituitary tumors, at 10 weeks post tamoxifen, neuroendocrine pituitary tumors were larger in the RPMax model (Fig. 2B). These data at the 10-week time point were consistent with long term survival analyses, where *Max*-deficiency led to highly significant reduction in time to morbidity (median 66 vs 162-day survival, p<0.0001) (Fig. 2C). Histological analyses revealed the presence of large pituitary adenomas and thyroid C-cell adenomas in both RP and RPMax models (Fig. 2D,E). Thus, *Max* loss strongly accelerates neuroendocrine pituitary and thyroid tumorigenesis in this sensitized mouse model. We previously showed that loss of *Max* in SCLC resulted in increased expression of proteins involved in serine biosynthesis and the one-carbon pathway (*32*). Western blot analyses showed that changes seen with MAX loss in SCLC such as an upregulation of SHMT1 and ATIC were also seen in the *Max*-deleted pituitary and thyroid cancers (Fig. 2F,G). We also examined features of different types of pituitary neuroendocrine tumors that arise in the RP, RPMax and Max-mutant alone (*Rb1/Trp53* wildtype) pituitary tumors using RNA-seq. Pituitary tumors in all three models expressed high levels of POMC, while we observed significant heterogeneity in expression of GH and Prolactin (Fig. 2H). Strikingly, pathways associated with MYC activity, such as one-carbon metabolism genes and ribosomal biosynthesis genes, exhibited increased expression in both the RPMax and the Max-mutant alone models compared to RP pituitary tumors, consistent with relative repression of these pathways by MAX (Fig. 2H).

**Figure 2:**
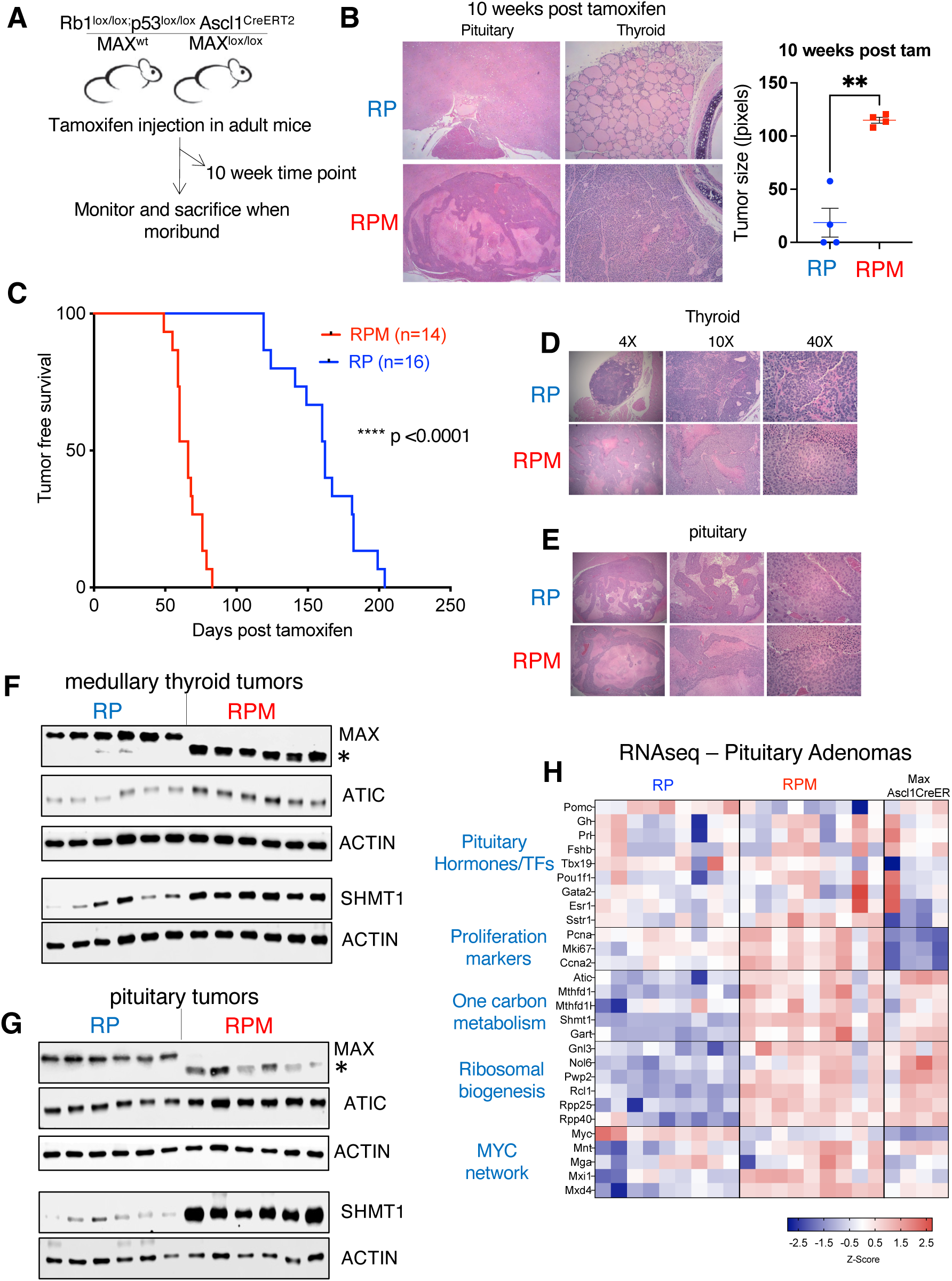
*Max* deletion cooperates with *Rb1/Trp53* loss to drive medullary thyroid carcinoma and pituitary adenomas. **A)** Illustration of mouse models employed to examine synergy between *Rb1/Trp53* and *Max* loss in neuroendocrine tumorigenesis. **B)** Hematoxylin and Eosin staining of pituitary and thyroid from RP vs RPMax models at 10-weeks post tamoxifen, with RP model exhibiting medullary hyperplasia and small pituitary adenomas while the RPMax model exhibiting bilateral medullary adenomas and large pituitary tumors. Quantification of increase in pituitary tumor size in the RPMax model shown to the right. **C)** Kaplan-Meier curve showing accelerated time to morbidity in the RPMax cohort with p-value fromlog-rank test shown. **D,E)** Mice in both RP and RPMax models exhibited medullary thyroid C cell and pituitary adenomas, with representative hematoxylin and eosin images of thyroid (D) and pituitary tumors (E) shown. **F,G**) Western blot analyses in thyroid tumors (F) and pituitary tumors (G) showing increased abundance of one carbon pathway proteins SHMT1 and ATIC with MAX loss. Asterisks indicate position of the N-terminal fragment of the truncated MAX protein in RPMax cells. **H)** Heat map showing RNA-seq data from pituitary tumors of RP and RPMax models as well as the *Max-*null only tumors (*Rb1/Trp53* wild-type) that arise with long latency.

### Differential gene expression in thyroid tumors and cell lines with wildtype and mutant MAX

To further characterize biological consequences and global alterations in gene expression due to *Max* inactivation, we derived cell lines from RP and RPMax primary thyroid tumors. These cells grew as clusters and spheres in suspension displaying a classical neuroendocrine phenotype as seen in SCLC. Western blot analyses revealed loss of the full-length MAX protein in cell lines derived from RPMax thyroid adenomas, with only the smaller N-terminal truncation band present in the mutant lines (Fig. 3A). Proliferation assays using CellTiter-Glo revealed increased growth in the four RPMax thyroid cell lines (Fig. 3B) compared to the RP-derived thyroid cancer lines, consistent with faster growth of thyroid tumors *in vivo* with *Max* loss. We restored MAX expression to a RPMax-derived thyroid cancer cell line H7536 using a doxycycline-inducible lentiviral vector and observed MAX protein increase (Fig. 3C) and suppression of proliferation as revealed by CellTiter-Glo experiments (Fig. 3D). These data confirm that MAX loss drives increased proliferation in this setting.

**Figure 3:**
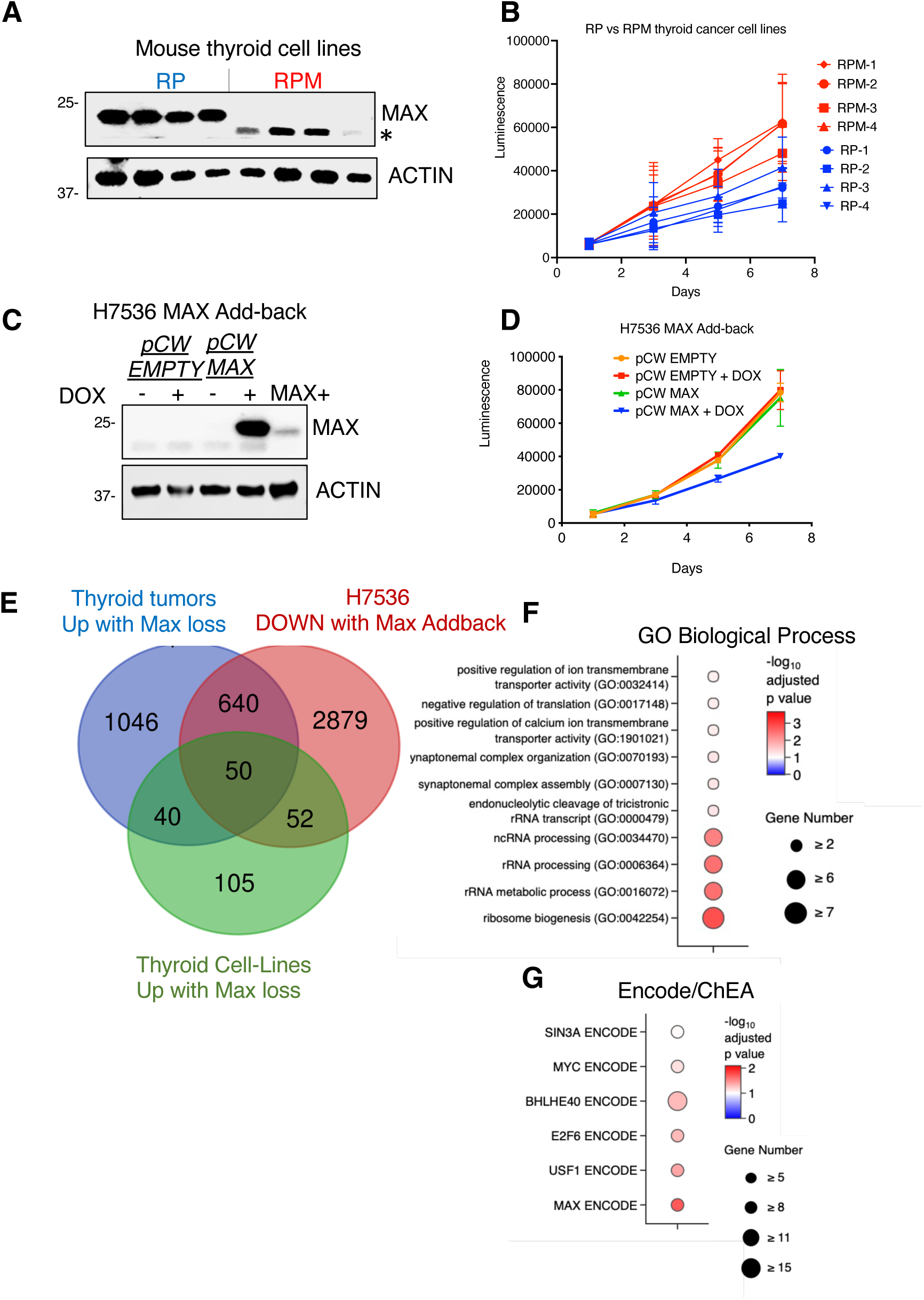
Increased proliferation in RPMax vs RP cell thyroid carcinoma cell lines. **A)** Western blot analysis showing loss of full-length MAX in thyroid cancer cell lines derived from the RPMax compared to those derived from the RP model. Asterisk indicates position of the N-terminal fragment of the truncated MAX protein in RPMax cells. **B)** CellTiter-Glo assays showing cell growth in 4 RPM and 4 RP thyroid cancer lines (n= 3 independent experiments). **C)** Western blot showing doxycycline-inducible return of MAX to an RPMax thyroid cancer cell line H7536. **D)** CellTiter-Glo assays showing suppressed cell growth upon return of MAX to the H7536 cell line (n=3 independent experiments). **E)** Venn diagram showing genes with increased expression upon MAX loss (EdgeR analyses, FDR <0.05) comparing RPMax to RP thyroid tumors, tumor-derived cell lines and genes with decreased expression upon doxycycline induced return of MAX to H7536. **F,G)** Pathway enrichment analyses querying GO Biological Processes (F) and ENCODE/CHEA datasets (G) including the 50 core MAX-regulated genes from (E).

To identify MAX-regulated genes in medullary thyroid cancer, we performed RNA-seq analyses. We compared gene expression in 6 RPMax vs 9 RP thyroid tumors, 4 RPMax vs 4 RP thyroid tumor-derived cell lines and in the RPMax-derived thyroid cancer cell line H7536 with inducible MAX re-expression (3 replicates per control vector vs *Max*-overexpressing condition). EdgeR analyses with a false discovery rate (FDR) cutoff of 0.05 revealed widespread changes in transcription upon MAX perturbation, with 3942 differentially expressed genes in the RPMax vs RP thyroid tumor comparison, 7569 in the MAX addback comparison and 602 in the RPMax vs RP tumor derived cell line comparison (Supplemental Tables 1-3). Our prior work in SCLC (*32*) highlighted tumor suppressor roles for MAX in gene repression, leading us to focus on genes upregulated upon MAX loss in thyroid tumors (1776 genes) and tumor-derived cell lines (247 genes), as well as genes downregulated upon return of MAX to RPMax H7536 cells (3622 genes). Overlap of genes upregulated in RPMax (compared to RP) tumors with those downregulated upon MAX addback to H7536 revealed 690 genes (Fig S1A). Pathway analyses using Enrichr (*40*) showed enrichment of MAX anti-correlated genes in ribosome biogenesis, translation and one-carbon metabolism (Fig. S1C/D). Including differential expression analyses of cell lines derived from RPMax vs RP tumors revealed a core set of 50 overlapping genes upregulated with *Max* deletion in tumors and tumor-derived cell lines and downregulated with MAX addback to H7536 cells. (Fig. 3E). Gene ontology analysis indicates that subsets of these core MAX-regulated genes function in ribosome biogenesis and translation (*Eif5a2*, *Rps2*, *Rrp12*, *Nol8*), synaptonemal complex formation and chromosome integrity (*Tdrd1*, *Stag3*, *Rad21*, *Syce2*, *Serf1*) as well as transcription (*Rorc*, *Hif1a*, *Taf4b*, *Mnt*) and metabolism (*Shmt1*, *Gart*) (Fig. 3F). Using ENCODE/ChEA motif analysis to identify classes of transcription factors likely associated with the differentially expressed genes underscores their relationship to regulation by MAX, MYC and bHLH proteins in general (Fig 3G, Fig S1B).

To delve more deeply into the relationship between gene expression and MAX loss of function, we assessed genomic occupancy of MAX in RP and RPMax thyroid tumor-derived cell lines using ChIP-Seq. While there was some variation in overall MAX occupancy at transcription start sites (TSS) in two cell lines derived from RP tumors, the RPMax thyroid cell lines showed strikingly diminished TSS MAX signal (Fig. 4A). A volcano plot of differentially expressed genes shows that over 10% of up-regulated, and 4% of down-regulated genes in RPMax lines were occupied at their transcriptional start sites by MAX in the MAX wildtype (RP) cells (Fig. 4B, blue dots). Moreover, genes occupied by MAX in RP cells were highly correlated with genes differentially expressed and up-regulated in RPM cells (Fig. 4 C). These differentially expressed genes include twelve of the fifty genes displaying overlapping MAX-dependent regulation in the tumors, the cell lines, and in MAX-addback cells (Fig. 4B, red dots; Fig. 3F). Tracks of individual genes exhibiting altered expression confirm loss of MAX occupancy at transcriptional start sites of loci differentially expressed in RPMax vs RP cells (Fig. 4D, showing MAX occupancy at Stag3 locus). Coverage plots showing MAX binding at transcription start sites in RP tumor cell lines indicate that loci upregulated in mutant MAX-derived RPMax cells corresponded to loci with the highest occupancy by MAX in RP cells. By contrast, downregulated genes in the RPMax cells corresponded to genes with lower MAX occupancy in RP loci (Fig. 4E).

**Figure 4.**
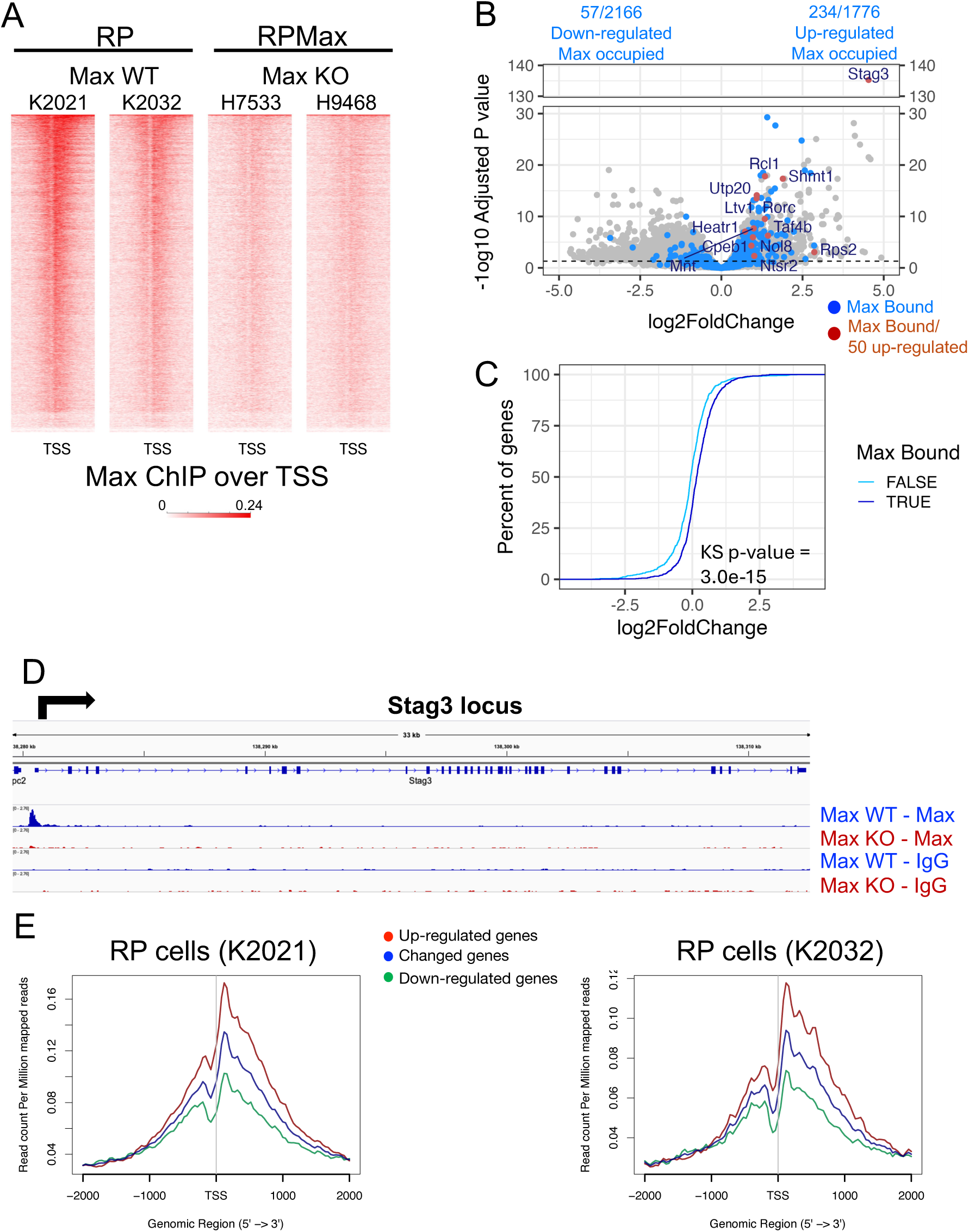
MAX genomic occupancy and correlation with gene expression changes in MAX-null thyroid tumors. **A)** Chromatin Immunopreciptation (ChIP) was performed on cell lines established from thyroid tumors with wild-type *Max* gene (RP, *Max* WT) or a mutant *Max* gene (RPMax, *Max* KO). Two of 3 biological replicates are shown. ChIP was carried out using anti-MAX or control IgG antibody. Heatmaps of MAX signal were generated centered on the transcription start site (TSS) +/- 2kb of flanking regions for each gene. **B)** Volcano plots of RNA-Seq data (comparing RPMax vs RP cell lines) were generated showing gene expression change (Log2FoldChange, x-axis) vs statistical significance (-log10 adjusted p value, y-axis) for every expressed gene comparing RPMax to RP cell lines. Genes occupied by Max ChIP in RP cell lines (identified by peakcalls) are shown in blue. Genes up-regulated in all data sets (50 genes, from Figure 3E) and bound by MAX are shown as red dots. The number of MAX bound genes that are up- and down-regulated are also shown. **C)** Cumulative distribution plots depicting the rank-order (y-axis, Percent of genes) of log2FoldChange (x-axis) of RNA-Seq data (comparing RPMax vs RP cell lines). MAX bound genes (dark blue) and genes determined to be MAX unbound (light blue) are shown. The difference between the distributions is statistically determined using Kolmogorov-Smirnov (KS) statistics. **D)** Genomic tracks showing Max occupancy at the *Stag3* gene locus in RP and RPMax cells. **E)** Line plots from ChIP experiments performed in (A) on Max WT RP cells were generated for all significantly changed (blue line), up-regulated (red), and down-regulated (green) genes in the RPMax cells. Each line is centered on the TSS (+/- 2kb) of every gene represented.

RPMax cells also display increased levels of RNA polymerase II (Pol2) on MAX occupied peaks (Fig. S2A), and at sites determined (by peakcalling) to be bound by MAX in RP cells (Fig. S2B), indicating considerable overlap between Pol2 and MAX occupancy. We note that Pol2 is generally increased in Max KO cells on gene loci that are differentially expressed upon MAX loss of function (Fig. S2B,C). This increase is observed at promoters, gene bodies, and termination regions (Fig. S2C). In addition, we were unable to detect a significant difference in RNA polymerase II travelling ratios i.e. the ratio of Pol II density in promoter relative to the gene body regions when we compared RPMax vs RP cells (Fig. S2D). This suggests that *MAX* inactivating mutations lead to a widespread increase in the accessibility of MAX bound loci to RNA polymerase II occupancy, irrespective of whether transcription is upregulated or downregulated.

### MAX inactivation leads to a shift in genomic binding by members of the proximal MYC network

MAX is an obligate heterodimeric partner for multiple members of the proximal MYC transcription factor network including the MYC protein family, the MXD family (MXD1-4, MNT) as well as MGA. Our RNA-seq data indicate a trend towards decreased expression of all three MYC paralogs in the RPMax compared to RP thyroid tumor cell lines (Fig. S3A). We and others had earlier reported that *MAX* inactivation resulted in accelerated degradation and loss of MYC protein (*17, 32–34, 41*). However, transcripts of other members of the MYC network (MNT, MGA, MXD1, MXD4, and MLX) are increased in the RPMax cells while MondoA (MLXIP) and ChREBP (MLXIPL) are largely unchanged (Fig. S3A). The MAX-like protein MLX, while unable to heterodimerize with MAX or MYC, forms heterodimers with the network members MondoA (MLXIP) and ChREBP (MLXIPL) as well as with MNT (*19, 21, 42*). Therefore, we asked whether loss of functional MAX, and subsequent downregulation of MYC, results in altered genomic binding by MNT, MondoA, and MLX in a manner that could affect survival and growth of the RPMax thyroid tumors.

We first assessed genomic binding by the MNT transcription factor. MNT is an HDAC-associated repressor that forms heterodimers with MAX and has been thought to antagonize MYC function (*42*). Nonetheless, MNT also promotes MYC oncogenic activity in B cell lymphomas through its suppression of MYC-induced apoptosis (*43*). MNT is also capable of dimerization with MLX where it functions as a transcriptional activator (*44*). Using peak calls from our ChIP-Seq data, we determined that MNT co-occupies approximately 30% of MAX-bound sites detected in RP cells (while also occupying a larger number of sites not bound by MAX) (Fig. S3B). By comparing MNT genomic occupancy in RP vs. RPMax tumor cell lines, we identified a cluster (denoted as MNT/MAX cluster 1) containing 84 MAX-bound gene loci that exhibited decreased MNT occupancy in the RPMax cells (Fig. S3C). Coverage plots, comparing overall occupancy over the genomic regions encompassing the 84 loci, demonstrate diminished association of both MAX and MNT at the transcription start sites of the RPMax relative to the RP tumor lines (Fig. 5A,B). Tracks at the *Rorc* locus are shown as a specific example (Fig. 5C). Integrating our genomic occupancy data with expression profiling indicates that these loci associated with MNT and MAX in RP cells are predominantly upregulated in RPMax cells, as shown in the volcano plot (Fig. 5D, red dots) and supported by cumulative distribution analysis, confirming the strong, statistical enrichment for genes upregulated upon Max loss (Fig. 5E). Enrichment analysis indicates that these 84 MNT/MAX cluster 1 genes were significantly comprised of MYC target genes, as well as loci involved in mTORC1 signaling, and the unfolded protein response (Fig. 5F). Therefore, a subset of genes encoding MYC targets are occupied, and at least partially repressed, by MNT-MAX dimers in RP cells. This MNT occupancy and transcriptional repression are lost when MAX is inactivated, resulting in upregulated gene expression in RPMax cells.

**Figure 5.**
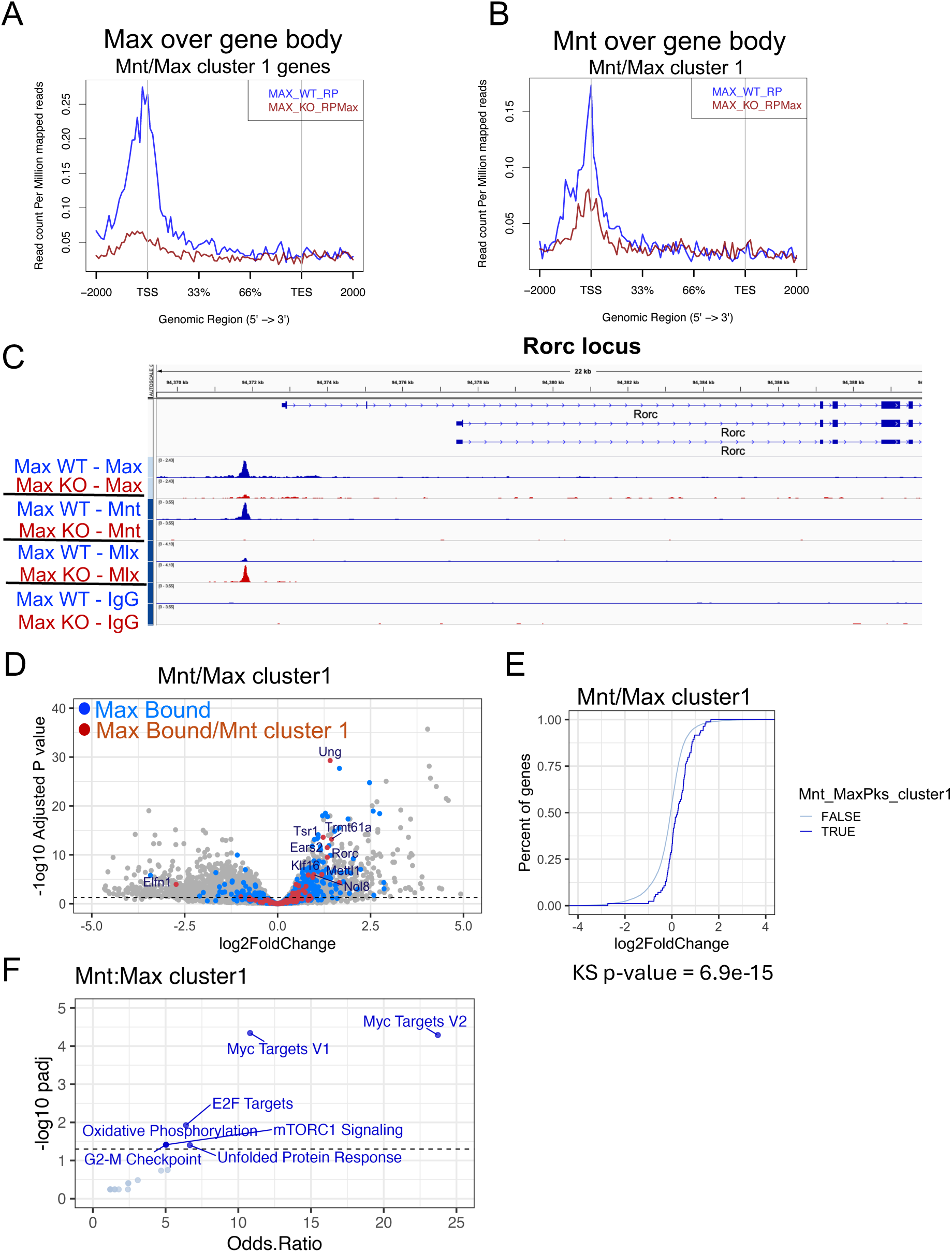
Alterations in MNT genomic occupancy correlates with gene expression changes in RP and RPMax tumor cells. **A)** Plots of genomic region over gene bodies (x-axis) vs normalized coverage (y-axis) of MAX/MNT bound cluster 1 (see Figure S3C), which was a cluster of peaks determined to be differentially occupied by MNT. Coverage of MAX for RP (MAX WT, blue) is compared to that from RPMax (MAX KO, red) cells. **B)** Plots of MNT coverage of MAX/MNT bound cluster 1 comparing RP (blue) and RPMax (red) cells, similar to panel A. **C)** Genomic tracks showing differential occupation of the *Rorc* promoter in RP (Max WT) and RPMax (Max KO) cells as determined by ChIP-Seq. The antibodies (against MAX, MNT, MLX) used for ChIP are shown (at left) after the hyphen. **D)** Volcano plot of RNA-Seq gene expression changes comparing RP vs RPMax thyroid tumors (Log2FoldChange on the x-axis, -Log10 adjusted p value on y-axis). Genes with peaks that map to MAX bound sites (blue) and in MAX/MNT cluster 1 (red) are shown. **E)** Cumulative distribution plots depicting the ranked-order (y-axis, Percent of genes) vs log2FoldChange (x-axis) of RNA-Seq data (comparing RPMax vs RP thyroid tumors). MNT cluster 1 genes (dark blue) and a similarly sized set of genes determined to not be MNT occupied (light blue) are shown. The difference between the distributions is statistically determined using Kolmogorov-Smirnov (KS) statistics. **F)** Enrichment analysis of genes found to be differentially occupied by MAX and MNT in Max WT RP tumor-derived thyroid cell lines (Mnt cluster1 from Fig. S5C). Enrichment categories are plotted as the -log10 of the adjusted P value (y-axis, -log10 padj), and Odds.Ratio (x-axis).

Prompted by previous studies showing that the MondoA (MLXIP) transcription factor is present as a heterodimer with MLX and influences MYC activity, we also assessed MondoA occupancy. To directly examine the effects of MAX on MondoA occupancy in the same cells, we used H7536 RPMax cells with doxycycline-inducible MAX vector. Heat maps show that treatment with doxycycline to drive MAX re-expression (MAX-addback cells) resulted in higher levels of MAX occupancy compared to controls, as expected (Fig. 6A). Importantly, we observed the opposite effect on MondoA occupancy, in that extensive association of MondoA with TSSs in RPM cells was strikingly diminished upon MAX-addback (Fig. 6A). Similarly, when we plotted MondoA peaks in RPMax cells over the corresponding MAX peaks (as detected in RP cells) we found that MondoA binding was decreased at these sites following MAX-addback (Fig. 6B). This suggests loss of MAX results in augmented MondoA binding. Further support for this conclusion comes from examination of gene tracks for the nucleophosmin gene, *Npm1* (Fig. 6C) and *Gart* (Fig. S5A). These genes are increased in expression with MAX loss and show reduced MAX binding and increased MondoA occupancy in the RPMax control tracks, with opposite effects in the MAX-addback cells. Thus, in the context of thyroid tumor derived cells, the absence of MAX correlates with replacement by MondoA at a subpopulation of loci whose expression is increased with MAX loss.

**Figure 6.**
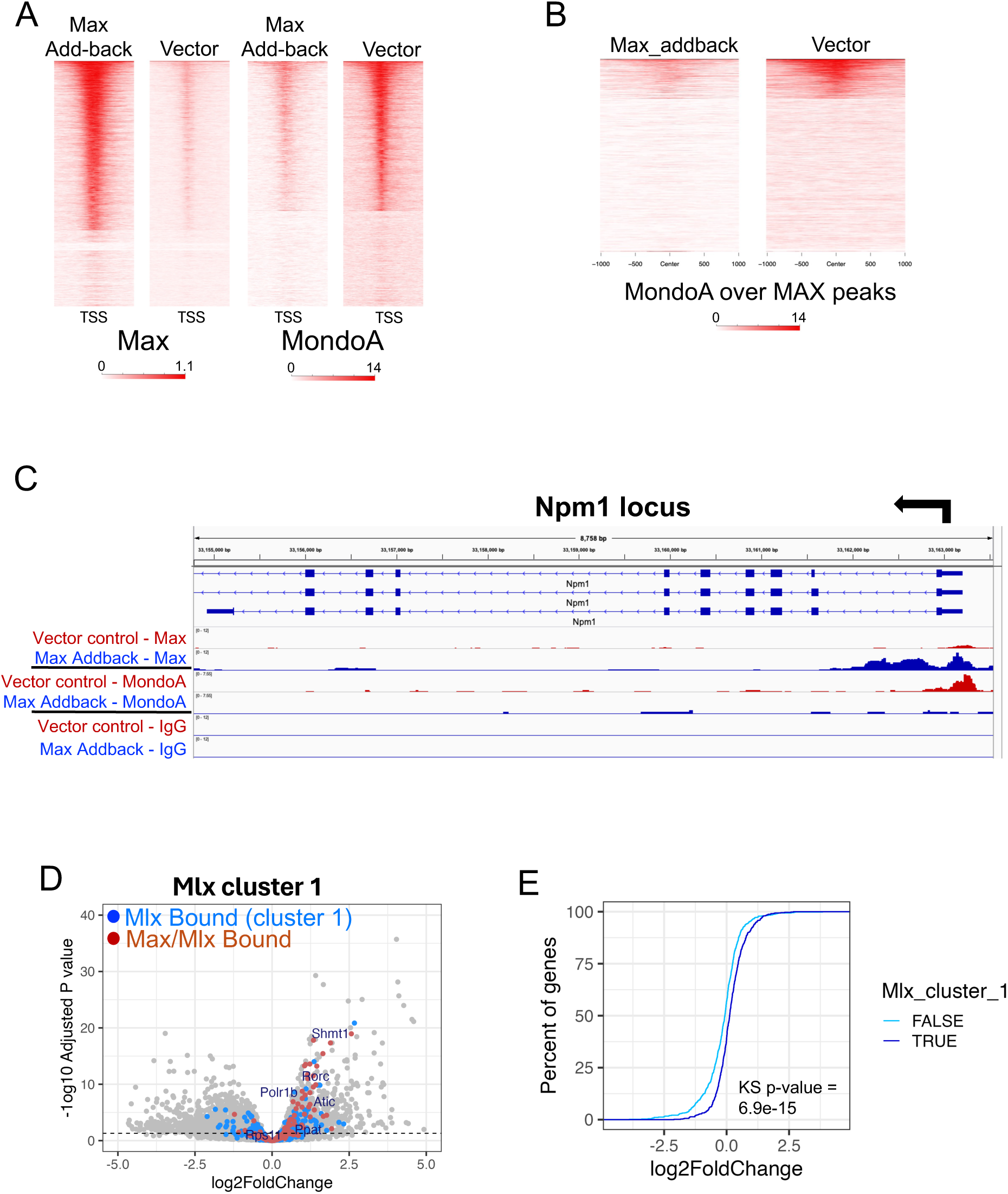
Altered MondoA genomic occupancy in RPMax tumor lines and MAX addback cell lines. **A,B)** Cut&Run heatmap analysis of MAX or MondoA in cells derived from *Max*-mutant thyroid tumor cells expressing DOX-inducible MAX (MAX-addback) or control vector (Vector). Cut&Run was performed on 1 million cells using the Auto Cut&Run robotic method with anti-Max, anti-MondoA, or control IgG antibody. Duplicate samples were merged, and heatmaps were plotted for Max (left) or MondoA (right) centered on the TSS +/- 2kb of flanking regions for every gene (panel A) or MondoA centered on MAX peaks as determined by peakcalls (panel B). **C)** Genomic tracks showing differential MondoA occupancy of the *Npm1* promoter in *Max*-null cells reconstituted with DOX induced MAX (MAX-addback), or vector control as determined by ChIP-Seq. The antibodies used for ChIP are shown (left) left after the hyphen for each cell type. **D)** Volcano plot of RNA-Seq gene expression changes comparing RP vs RPMax thyroid tumors (Log2FoldChange on the x-axis, -Log10 adjusted p value on y-axis). Genes with peaks that map to Mlx cluster 1 (blue) and both Mlx cluster 1 and Max (red) are shown. **E)** Cumulative distribution plots depicting the rank-order (y-axis, Percent of genes) vs log2FoldChange (x-axis) of RNA-Seq data (comparing RPMax vs RP thyroid tumors). Mlx cluster 1 genes (dark blue) and a similar sized set of genes not in an Mlx occupied cluster (light blue) are shown. The difference between the distributions are statistically determined using Kolmogorov-Smirnov (KS) stats.

Because MLX dimerizes with both MondoA and MNT (*19, 44*), we determined the genomic occupancy of MLX in RPMax cells. Plotting MLX coverage over all transcription start sites revealed a set of genomic loci (denoted as MLX cluster 1) with increased MLX binding in the absence of MAX (Fig. S5B, C). This cluster contains genomic loci that are strongly enriched for up-regulated genes in *Max* mutant tumor cells (Fig. 4D,E). Importantly, this cluster also contained 22 of the 84 genes previously found to lose MAX/MNT binding upon *Max* deletion (Fig. 5A-F), indicating that MLX likely occupies promoters for many of these up-regulated genes in the absence of MAX (e.g., the *Rorc* locus shown in Fig 5C).

Surprisingly, we noted significant de novo MAX- and MLX-independent genomic occupancy by both MNT and MondoA in RPMax cells (Fig. S3B, Fig. S5D). To further investigate how MAX loss affects MNT binding, MNT coverage was plotted and clustered over transcription start sites (TSS) of all genes (Fig. S3C). One MNT cluster occupied regions distal to the TSS, and gene bodies (denoted as MNT TSS cluster; Fig. S3D). We further filtered this MNT occupied cluster of genes based on the approximately 15% that were also MAX bound in RP cells. MAX binding to this cluster of genes was centered near the TSS, and this was abrogated in RPMax cells (Fig. S4A). MNT occupancy, however, was not decreased in the MAX bound gene bodies and TSS proximal regions of this cluster (Fig. S4B). Importantly, these genes, which include the MNT locus itself, were strongly de-repressed in the RPMax cells (Fig. S4C-E). This cluster was enriched for genes involved in cell cycle and mitosis (Fig. S4F) but did not include MYC target genes. Similarly, MondoA binding in *Max*-deleted cells, while overlapping WT MAX sites, appears to be largely independent of MLX binding (Fig. S5D). Therefore, both MNT and MondoA possess the capacity to occupy sites unoccupied by either MAX or MLX, perhaps through homodimerization or with an as yet unidentified heterodimeric partner. These data suggest a significant reshuffling of heterodimeric partners and binding sites in Max-deleted tumor cells resulting in expression changes in multiple genes.

### Differential sensitivity of MAX-inactivated thyroid tumors to inhibition of MNT, MLX and MondoA

Given the observed changes in genomic occupancy of MYC network members MNT, MLX, and MondoA in thyroid tumors with WT vs *Max* mutant alleles, we next assessed whether genetic or pharmacologic suppression of these factors impacts growth of the RP or RPMax tumor cells. We therefore determined the number of colonies appearing after low-density plating subsequent to introduction of siRNAs against MLX or MNT into RP and RPMax lines. Figure 7A shows the relative fraction of colonies per well produced by three independent RP and RPMax cell lines following siRNA treatment. The specificity of the siRNAs employed for this study have been previously validated in several different biological settings (*24, 45*) and the strong effect of the siDeath indicates adequate transfection. The data indicate that siMnt has a modest and variable suppressive effect on growth of the RP tumor cells which is further suppressed in the *Max*-deleted RPMax cells (Fig. 7A). By contrast, siMLX had little effect on growth of the RP lines but significantly attenuates growth of the RPMax cells (Fig. 7A). The sensitivity of RPMax cells to MLX knockdown prompted us to employ a small molecule probe (SBI-477), previously derived from a high-throughput screen as an inhibitor of MondoA that suppresses expression of MondoA’s key downstream targets (*46*). We therefore determined the effects of 20μM SBI-477 on RP and RPMax tumor cells, a concentration we determined to inhibit growth of other MondoA-dependent tumor lines. Here we find that, relative to growth on untreated cells, SBI-477 treatment had no significant effect on proliferation of RP cells while it selectively inhibited growth of RPMax cells (Fig. 7B). Lastly, we tested the effect of SBI-477 on H7536 RPM cells in which MAX was reintroduced using the doxycycline-inducible system (Max-addback cells) resulting in a reduced MondoA genomic occupancy (Fig. 6A,B). As expected, proliferation of RPMax cells infected with vector controls (pCW vector), with or without doxycycline, were sensitive to SBI-477 as were the uninduced MAX-addback cells (pCW MAX). However, in the presence of doxycycline-induced MAX, sensitivity to SBI-477 was significantly reduced (Fig. 7C). These data support the idea that loss of MAX leads to increased dependence on MondoA function in these thyroid tumor cells and are consistent with our findings that MondoA proteins occupy a subset of loci that are normally MAX targets in wildtype tumors and are differentially expressed in RPMax tumors.

**Figure 7.**
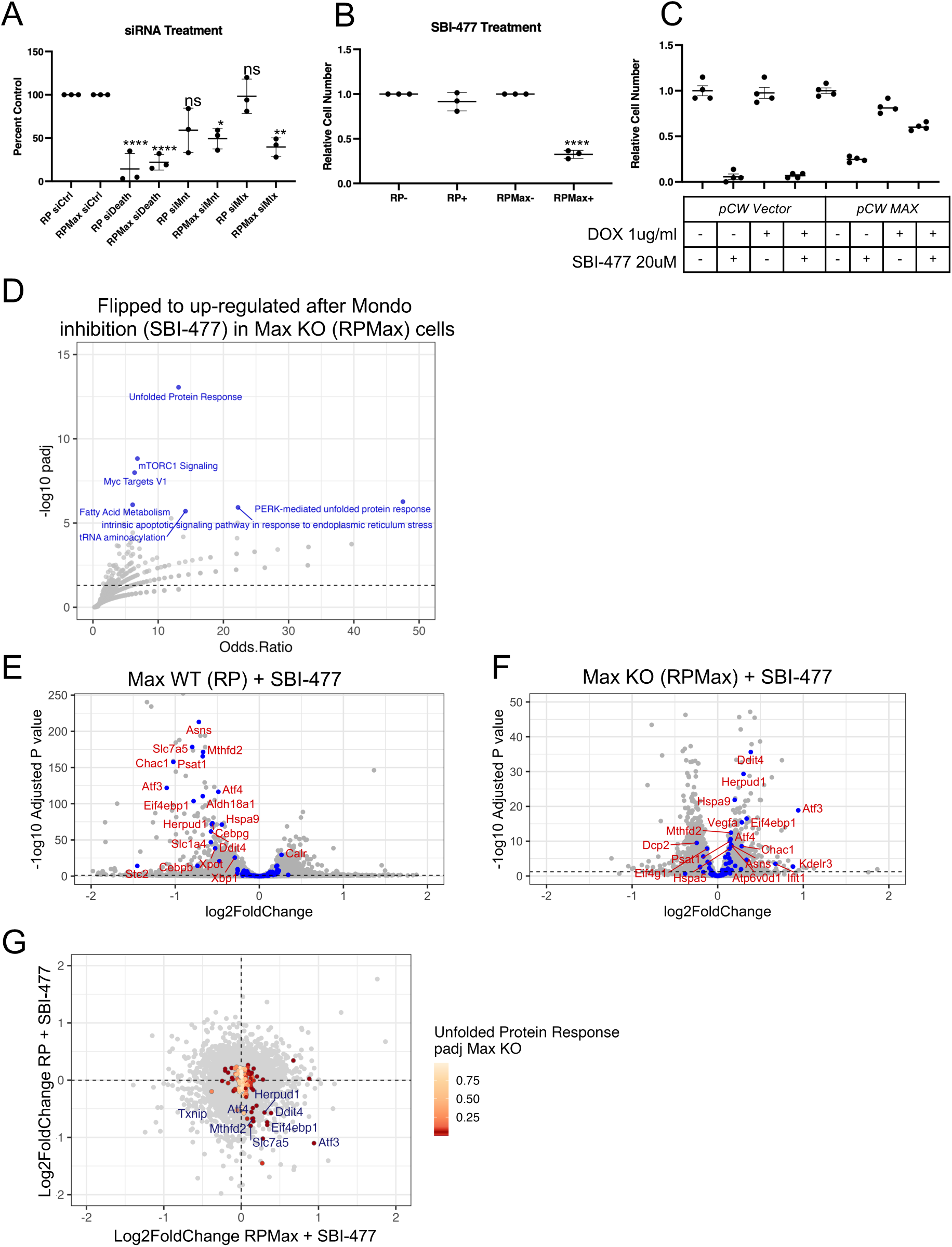
Differential sensitivity and changes in gene expression of *Max*-inactivated thyroid tumor upon inhibition of MNT, MLX and MondoA. **A)** Thyroid tumor cell lines (3 independent RP and 3 RPMax lines) were transfected with the indicated siRNA and allowed to reconstitute over 4 days. Outgrown spheres were enumerated and normalized to the siCtrl condition. siDeath is included as a control for transfection efficacy. **B)** Thyroid tumor cell lines (3 independent RP and 3 RPMax lines) were seeded at equal density with either vehicle (DMSO) or SBI-477 (Mondo Inhibitor, 20uM). After 4 days, cells were counted and normalized to DMSO. **C)** RPMax thyroid tumor cell line H7536, reconstituted with either control vector (pCW Vector) or doxycycline-inducible Max construct (pCW MAX) were cultured in the absence or presence of DOX and/or SBI-477 for 96 hours then cells were counted and normalized to untreated cells for each line. For all data, p-values were calculated using ANOVA with a Tukey’s adjustment. **D)** Enrichment analysis of genes that were down-regulated in tumor cells expressing wild-type *Max*, but up-regulated in tumor cells with *Max* deleted upon MondoA silencing (using a Mondo inhibitor, SBI-477). Enrichment categories are plotted as the -log10 of the adjusted P value (y-axis, -log10 padj), and Odds.Ratio (x-axis). **E, F)** Volcano plots of RNA-Seq gene expression changes (Log2FoldChange on the x-axis, -Log10 adjusted p value on y-axis) comparing cells treated with vehicle control to cells treated with MondoA inhibitor (SBI-477) for RP cells (D) or RPMax cells (E). Genes that are important in the unfolded protein response are highlighted in blue. **G)** Plot of expression changes (Log2FoldChange) of RP (y-axis) vs RPMax (x-axis) cells. Genes important for the unfolded protein response are highlighted by yellow-red color. Yellow-red shading depicts the adjusted p value (padj), and higher red intensity indicates greater significance in gene expression change between the two datasets.

To explore the consequences of SBI-477 treatment in more detail we carried out RNA-Seq on RP and RPMax cells to determine changes in RNA expression in mock-treated and drug-treated cells. When we focused on expression changes in loci upregulated in RPMax cells following SBI-477 treatment, we detected a highly significant increase in transcripts related to stress response such as mTORC1 signaling and particularly, the unfolded protein response (Fig. 7D). Many of the same genes and other related genes were downregulated in RP cells subsequent to SBI-477 treatment (Fig.7 E,F). Among the most affected were *Atf3*, *Atf4*, *Ddit4*, *Herpud*, *Chac1* all encoding factors linked directly or indirectly to the ER stress response (Fig. 7E,G). These data reveal significant differences in molecular responses to MondoA pathway inhibition in the RP and RPMax cells suggesting that MondoA acts in the RPMax tumor cells to suppress the unfolded protein response which results in cell death.

## Discussion

Here we report findings that confirm and expand the role of MAX as a neuroendocrine tumor suppressor. We demonstrate that *Max* deletion alone in mice is sufficient to induce pituitary adenomas that arise with long latency. This is significant, as this tumor type arises in a subset of patients harboring germline mutations in *MAX* (*28*). The long latency suggests a requirement for multiple oncogenic events that cooperate with *Max* inactivation in order for tumors to evolve. Moreover, we demonstrate in *Trp53* and *Rb1* mutant mice (RP) that *Max* inactivation (RPMax) dramatically accelerates thyroid and pituitary tumorigenesis. This work builds on earlier observations indicating that *MAX* undergoes loss of function mutations in multiple human neuroendocrine cancer types including pheochromocytoma and pituitary adenoma as well as SCLC and other cancer types (*26–28, 30, 31, 36, 47*). Recently we found *Max* deletion as a top hit in a genome-wide CRISPR/Cas9 screen designed to identify functional tumor suppressors in preneoplastic lung neuroendocrine progenitor cells (preSC) (*32*). In addition, we reported a dramatic acceleration of SCLC in RP mice with targeted inactivation of *Max* in lung epithelium (*32*). In the present study we employ an inducible Cre recombinase under control of the *Ascl1* promoter in order to delete *Max* in neuroendocrine cells of adult RP mice. While both RP and RPMax mice developed C-cell (medullary) thyroid tumors and pituitary adenomas, the RPMax mice displayed significantly increased tumor size at ten weeks and an approximately 60% decrease in median survival time relative to the RP mice. These data, together with the previous studies, including MAX germline mutant patient phenotypes (reference Seabrook 2021 ref) establish *MAX* inactivation as an initiator of multiple endocrine neoplasia (MEN) syndrome.

Homozygous *Max* loss of function appears to take multiple forms in MEN. The initial description of *Max* loss in PC12 cells indicated intragenic deletion of exon 4, thereby abolishing the ability of MAX to homo-or heterodimerize (*25*). Analysis of *MAX* loss in SCLC tumors and cell lines demonstrated intragenic deletions often spanning multiple exons (*29*). In addition, several reports indicate scattered point mutations leading to premature translational terminations or exon skipping (*27, 30, 31*). In the cases where it was examined, no MAX protein was detectable using a C-terminal MAX antibody (*29*). In our study we generated a targeted deletion of 841bp encompassing *Max* exon 4 resulting in deletion of the helix-2-leucine zipper domain thereby producing a reading-frameshift and translational termination in non-coding exon 5 (*17*). While a C-terminal anti-MAX was unable to detect any full-length MAX protein (*17, 32*), we observed that an N-terminal anti-MAX detected a lower molecular weight form (14kDa), consistent with the predicted frameshifted termination product of 127 amino acids. As expected, this truncated protein was not observed in the completely deleted CRISPR/Cas9-*Max* in preSC cells which consistently exhibited increased proliferation and survival relative to preSC cells expressing full-length functional MAX (*32*). Interestingly, it was recently reported that the intrinsically disordered N-terminal 21 residue segment of MAX is capable of mediating increased on- and off-rates of MYC-MAX heterodimer binding to E-box DNA, thereby promoting selective binding to high-affinity E-box sites (*48*). We think it highly unlikely that this N-terminal MAX segment plays a role in the neuroendocrine tumorigenesis reported here, since MYC is rapidly degraded and undetectable in the *Max* null cells, including RPMax cells, and moreover, the same targeted *Max* allele is unable to promote MYC dependent oncogenesis in the contexts of either lung epithelial cells or Eμ-*Myc* transgenic mice (*17, 32*).

The role of *MAX* inactivation in MEN poses an apparent paradox-namely, that the obligate dimerization partner for the MYC oncoprotein, a major driver of a broad spectrum of cancers, nonetheless acts as a tumor suppressor. We surmise that this dual activity of MAX lies in its ability to dimerize, not only with MYC, but with other members of the MYC network (*1*). MAX heterodimerizes with the MXD protein family (MXD1-4), and the MNT and MGA proteins (*4, 42, 49–51*). All these proteins can function as transcriptional repressors, likely playing roles in growth arrest and differentiation, and several have been shown to act as tumor suppressors (*52–54*). Inactivation of *MAX* would be expected to abolish or attenuate the activities of these factors in addition to causing loss of MYC function. Such a drastic “disconnect” of essential regulatory proteins within the MYC network might be anticipated to induce stress and apoptosis. Yet the MLX protein, while unable to substitute for MAX in binding to MYC, MGA or most MXD proteins, has been shown to be capable of dimerization with MNT as well as with MondoA (MLXIP) and ChREBP (MLXIPL) (*21*) (*20*) (*19*).

Considering the correlation observed between the levels of MAX promoter occupancy in the RP tumor cells and RNA polymerase binding and altered gene expression in the RPMax cells, we focused on occupancy of network proteins in RPMax cells specifically at sites associated with MAX binding in RP cells. Taken together our data indicate that the predominant transcriptional programs differentially expressed in the *Max*-null tumor cells are linked to genes that are directly bound by MNT, MLX or MondoA and include MYC target genes, cell cycle checkpoints, mitotic, and E2F targets as well as GART, a key regulator of purine biosynthesis. Furthermore, because MNT knockdown (Mlx knockdown as well) or treatment with an inhibitor of the MondoA pathway induces distinct transcriptional responses and preferentially blocks growth of *Max*-null relative to *Max*-wildtype tumor cells, we conclude that as a consequence of *Max* inactivation, the altered binding of MLX, MondoA and MNT are likely to be critical for the survival of RPMax cells.

Based on the above, we propose a scenario linking MAX loss to tumorigenesis (see Fig. 8). While MYC-MAX heterodimers are widely essential for cell proliferation, MYC abundance can vary significantly in different cell types and cell states. For example, MYC levels are usually low to undetectable in many primary, as well as stem and progenitor cells relative to tumors with dysregulated MYC (*55*). Moreover, normal B lymphoid cell survival and differentiation can be initiated in the absence of endogenous MYC, MAX or both (*17, 56*). Even tumor cell populations exhibit pulsatile and heterogeneous MYC expression (*57*). Cells passing through a transient state in which MYC levels are low would be better capable of tolerating MAX loss. Furthermore, with the accompanying inactivation of growth inhibitory MXD proteins, *Max* mutant cells may transiently maintain critical MYC target gene expression and acquire an augmented capacity for proliferation and survival. The absence of wildtype *Rb1* and *Trp53* would likely promote or contribute to this survival and growth. Such cells might eventually senesce or die as a result of failure to respond to proliferative signals. However, a subpopulation able to transition to augmented growth mediated by the significantly altered genomic occupancies of Mnt, MLX and MondoA could in principle bypass the requirement for MYC and, in addition, exhibit an abnormal or discordant response to mitogenic or growth signals that would lead to cell-autonomous growth and eventually neoplasia (Fig. 8). Of course, it is also plausible that other transcription factors, particularly of the bHLH E-Box binding class, would contribute, directly or indirectly, to the rescue and transformation of MAX loss-of-function cells. Transcription factors such as MGA, MITF, TFEB, ASCL1 and Clock proteins are strong candidates. Genomic binding by the E2F family of transcription factors has also been associated with MYC-MAX occupancy and these factors could potentially contribute to gene regulation in the absence of MAX (*58–60*). It will be important in the future to determine the nature and dynamics of the factors that control critical loci during neuroendocrine tumor progression. Nonetheless, our study indicates that perturbations of the MYC network can provide insights into the transcriptional mechanisms that underlie cell plasticity.

**Figure 8.**
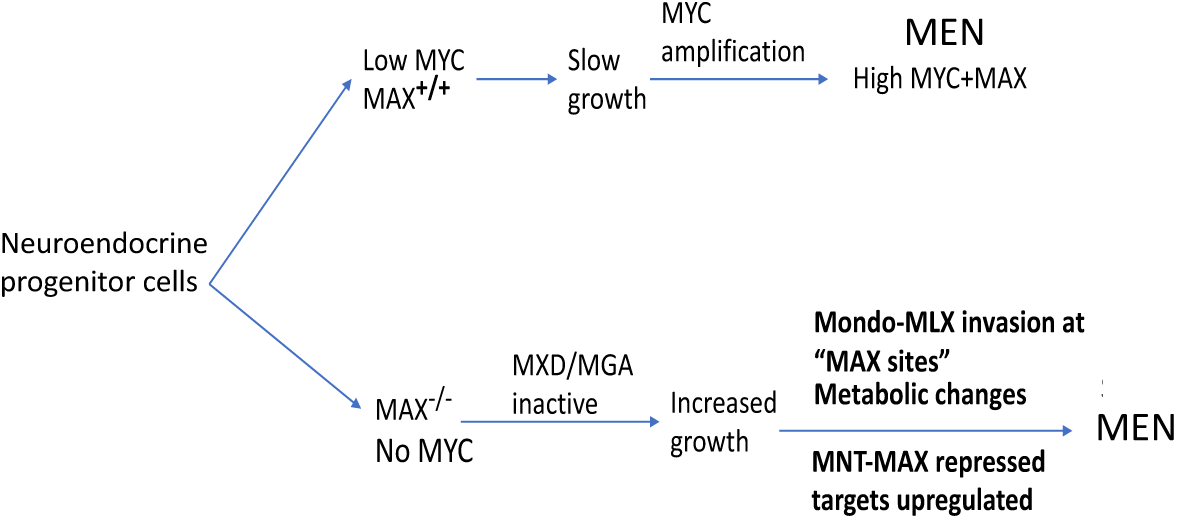
Diagram showing predicted stages in the development of tumors derived from cells with inactivated *MAX* (see Discussion).

## Materials and Methods

### Mouse models

The *Rb1*^lox/lox^ strain was originally obtained from Dr. Tyler Jacks, the *Trp53*^lox/lox^ strain from Dr. Anton Berns, and the *Ascl1-Cre-ERT2* strain from Dr. Jane Johnson. We bred *Ascl1-Cre-ERT2* and compound mutant *Rb1*^lox/lox^;*Trp53*^lox/lox^;*Ascl1-Cre-ERT2* mice to a *Max*^lox/lox^ allele (*32*). Tamoxifen induction of Cre-ERT2 under the control of *Ascl1* promoter was accomplished by i.p. injection of mice with 150 mg/kg/day tamoxifen, prepared in sterile corn oil, over 5 consecutive days. Mice were euthanized when they exhibited poor body condition and/or hunched appearance, or when flagged with visible thyroid tumors. Necropsies were performed and inner organs fixed in 10% neutral buffered formalin (NBF) or, for bony tissues and the skull, Bouin’s fixative. Tissues were fixed for 24 hours (NBF) or 1 week (Bouins) before paraffin embedding. Tissue sections (4 μM thick) were stained with Hematoxylin and Eosin (H&E). Upon necropsy, any tumors present tumors were split with one portion used for fixation and histological analyses, a second portion was snap frozen for molecular analyses and, for a subset of tumors, cell lines were generated by breaking up tumors mechanically and plating in cell culture media. All animal procedures were approved by the Institutional Animal Care and Use Committee (IACUC) at the Fred Hutchinson Cancer Center.

### Cell lines

Mouse cell lines were extracted as mentioned above and grown in DMEM (11965-092, ThermoFisherScientific) supplemented with penicillin (100 U/ml), streptomycin (100 μg/ml) (15140122, ThermoFisherScientific), and 10 % fetal bovine serum (FB-01, Omega Scientific). Cells were maintained at 37°C in a humidified atmosphere containing 5 % CO2 and 95 % air.

### Cell Proliferation and Viability

To monitor cell proliferation, 2,000 cells were seeded per well in 96-well plates. Each condition was tested with 3 independent replicates, and each replicate included 4 wells. The cells were incubated for 7 days. Cell viability was assessed using the CellTiter-Glo assay (Promega) according to the manufacturer’s instructions. Data were analyzed and visualized using GraphPad Prism software.

### Western Blot Analysis

Whole-cell protein extracts were prepared in cold cell lysis buffer (20 mM Tris-HCl (pH 7.5), 150 mM NaCl, 1 mM Na2EDTA, 1 mM EGTA, 1 % NP-40, 1 % sodium deoxycholate) supplemented with protease and phosphatase inhibitors (78441, ThermoFisherScientific). Tumor samples were homogenized in cold cell lysis buffer using the AgileGrinder™ Tissue Grinder (ACT-AG3080, Thomas Scientific-ACTGene). Proteins were quantified using Pierce™ BCA Protein Assay Kit (23227, ThermoFisherScientific), resolved on 4–20 % Mini-PROTEAN®TGX™ Precast Protein Gels (4561096, BioRad), and transferred to Amersham Protran 0.45 NC nitrocellulose membranes (10600002, GE healthcare life science). Protein samples were normalized to ACTB or HSP90. Imaging was performed using the LI-COR Odyssey Fc.

### Lentiviral infections

Lentiviral transduction of H7536 thyroid cancer cells was performed as described (*32*) to express MAX ORF Isoform 2 (NM_145112.2) that was cloned into pCW57-RFP-P2A-MCS (Addgene plasmid no. 78933).

### Transfection of siRNA into thyroid tumor lines

Suspension cell line spheres were pelleted, washed with PBS and disassociated with Accutase solution (Sigma-Aldrich) for 10 minutes at 37°C to single-cell suspension. Cell suspension was then transfected with siRNA using RNAi-Max reagent (Invitrogen) and Opti-MEM (Gibco). The siRNA used were Flexitube siRNA (Qiagen), a combination of 4 unique siRNAs per target, as well as All-Star non-targeting negative control and All-Star Cell Death siRNA, as a positive control for cell killing, and thus a proxy for transfection efficiency (Qiagen catalog numbers; siMnt: GS17428, siMlx: GS21428, All-Star Negative Control: 1027280 and All-Star Cell Death: SI04939025). Transfected cells were monitored for up to 4 days for sphere-formation and viability.

### SBI-477 treatment of thyroid tumor lines

The MondoA inhibitor SBI-477 (MedChemExpress) was reconstituted in cell culture grade dimethyl-sulfate, DMSO (ChemCruz), as a 10mM stock. Thyroid tumor cell lines were treated with either vehicle alone (DMSO) or 20uM SBI-477. Treated cells were monitored for viability. For RNA-Seq, triplicate samples were DMSO versus SBI-477 treated for 72 hours, then pelleted for harvesting. RNA was harvested using Direct-Zol RNA minipreps (Zymo Research) following the manufacturer’s recommendation. Samples were submitted to the Fred Hutch sequencing core for Tru-Seq mRNA sequencing.

### RNA-Seq analysis

RNA was extracted using TRIZOL according to the manufacturer’s recommendations (15596018, ThermoFisherScientific). RNA-seq libraries prepared using the Ultra RNA Library Prep Kit for Illumina (E7530L, New England BioLabs) from total RNA (500 ng). All library preparation was conducted according to the manufacturer’s instructions. Single-end sequencing (50 bp) was performed using an Illumina HiSeq 2500, and reads of low quality were filtered before alignment to the mm9 genome build using TopHat v2.0.12 (*61*). Cuffdiff v2.1.1 (*62*) was used to generate FPKM expression values.

### ChIP-Seq and Cut&Run

Chromatin immunoprecipitation and sequencing (ChIP-seq) was done for MAX, MNT, and RNA-Pol2 using an MNase digestion step to allow nucleosome resolution of ChIP fragments (*63*). Briefly, after formaldehyde cross-linking, cell lysis, and chromatin fragmentation with MNase, the final SDS concentration after dilution of total chromatin was increased to 0.25% with addition of 20% SDS stock solution. Sonication was performed in a Covaris M220 focused ultrasonicator for 12 minutes with the following settings: 10% duty cycle, 75W peak incident power, 200 cycles/burst, and 6-7°C bath temperature. The SDS concentration of the sonicated chromatin solution was readjusted to 0.1% with dilution buffer. Immunoprecipitation was performed using antibodies against RNA Pol2 (4H8, Active Motif), MAX (10426-AP from Proteintech, or C17 from Santa Cruz Biotechnology), MNT (A303-627A, Bethyl Laboratories), MLX (D8G6W, Cell Signaling Technology) or negative control IgG (Cell Signaling Technology) using 10 μg of antibody for each immunoprecipitation on the clarified chromatin (input) fraction from 10×10^6^ cellular equivalents. DNA was then purified using standard phenol:chloroform extraction, 10 pg. of spike-in DNA purified from MNase-digested chromatin from *S. cerevisiae* was added to permit comparison between samples. Single strand library preparation for ChIP samples was then performed as described (*64*).

Samples were then submitted for 25×25 paired-end sequencing (5-10 million reads for CUT&RUN, and 20-30 million reads for ChIP) on an Illumina HiSeq 2500 instrument at the Fred Hutchinson Cancer Research Center Genomics Shared Resource.

Genomic binding sites for MAX, and MONDOA were determined using Cleavage Under Targets and Release Using Nuclease (CUT&RUN) using the Auto CUT&RUN system at the Fred Hutch Cancer Center (*65, 66*). Briefly, CUT&RUN was performed on cells derived from thyroid tumors with normal *Max* gene (RP cells) or mutant *Max* (RPMax cells). The spheroid cells were treated with Accutase to achieve single cell suspension, bound to ConA beads, then permeabilized, and incubated overnight with antibody to either MAX (10426-AP from Proteintech, or C17 from Santa Cruz biotech) or MONDOA/MLXIP (13614-1-AP, Proteintech). Duplicate samples were performed for each antibody used. After digestion with secondary bound protein-A MNase (Henikoff lab, FHCC), the reaction was quenched, and 2 ng of yeast ‘spike-in’ DNA was added. DNA fragments released into the supernatant were collected and directly added without purification into an end repair and dA tailing reaction done at 58°C (to enhance the capture of small DNA fragments). TruSeq adapters were ligated (Rapid DNA ligase, Enzymatics) onto the DNA, and DNA was digested with Proteinase K. DNA was then size fractionated using Ampure Beads (Beckman-Coulter) and library preparation was then carried out (KAPA Biosystems) by adding barcoded primers. The size distribution was determined for each sample using an Agilent 4200 TapeStation, and quantity of DNA was determined using Qubit (Invitrogen).

Sequences were aligned to the mm10 reference genome assembly using Bowtie2 with the arguments: --end-to-end --very-sensitive --no-mixed --no-discordant --overlap --dovetail -I 10 -X 700. Datasets were also aligned to the sc3 *(S. cerevisiae*) assemblies to enumerate reads from spike-in DNA.

Counts per base-pair were normalized as previously described by multiplying the fraction of mapped reads spanning each position in the genome-by-genome size or by scaling to spike-in DNA. Peak calling was done using the MACS2 package. Peaks were considered to be associated with a gene if the peak was present in the promoter or genebody.

### Computational analysis

Downstream analysis of differentially expressed genes by RNA-Seq and correlation with MAX, MNT, MLX, MONDOA, and RNA pol2 occupation, with CUT&RUN and ChIP-Seq data, was carried out using custom R scripts and functions developed in the Eisenman lab. The GenomicRanges R package was used to process genome positions. Genomic plots were made using ngs.plot (https://github.com/shenlab-sinai/ngsplot) or the R package ggplot2. Clustering was performed by plotting all genomic regions centered on the TSS using 5 k-means clusters for each sample (using ngs.plot). Bedtools was also used to numerate genomic positions. Duplicate samples for CUT&RUN analysis were merged together for increased coverage, and peakcalls were done on the merged files. Application of Kolmogorov-Smirnov (KS) statistics was used to assess the significance of cumulative distribution plots by assessing genes of interest (MAX, MNT, MONDOA, or MLX occupied) compared to the same number of a random set of expressed genes. Traveling ratios were assessed from RNA-Pol2 reads derived from ChIP-Seq data by dividing promoter reads by genebody reads for each gene.

## Acknowledgements

We are grateful to Pei-Feng Cheng for generating the Max mutant mice and to Xiaoying Wu and Yulong Su for critical comments on the manuscript. The authors acknowledge the contributions of Fred Hutch Shared Resources, including Comparative Medicine (RRID:SCR_022610), Experimental Histopathology, Genomics and Bioinformatics of the FredHutch/University of Washington/Seattle Children’s Cancer Consortium (P30 CA015704). Scientific computing infrastructure was supported by the Office of Research Infrastructure Programs (S10OD028685). This work was supported by National Institutes of Health (NIH) National Cancer Institute grants RO1 CA248762 (to R.N.E. and D.M.) and R35 CA231989 (to R.N.E.)

## Author contributions

Conceptualization: DM, RNE

Methodology: DM, AI, BF, AA

Investigation: BF, AI

Visualization: BF, AI, AA

Supervision: DM, RNE, AA

Writing—original draft: DM, RNE

Writing—review & editing: All Authors

