## Supplementary Figures + Legends for "MAX inactivation deregulates the MYC network and induces neuroendocrine neoplasia in multiple tissues"

Fig.S1

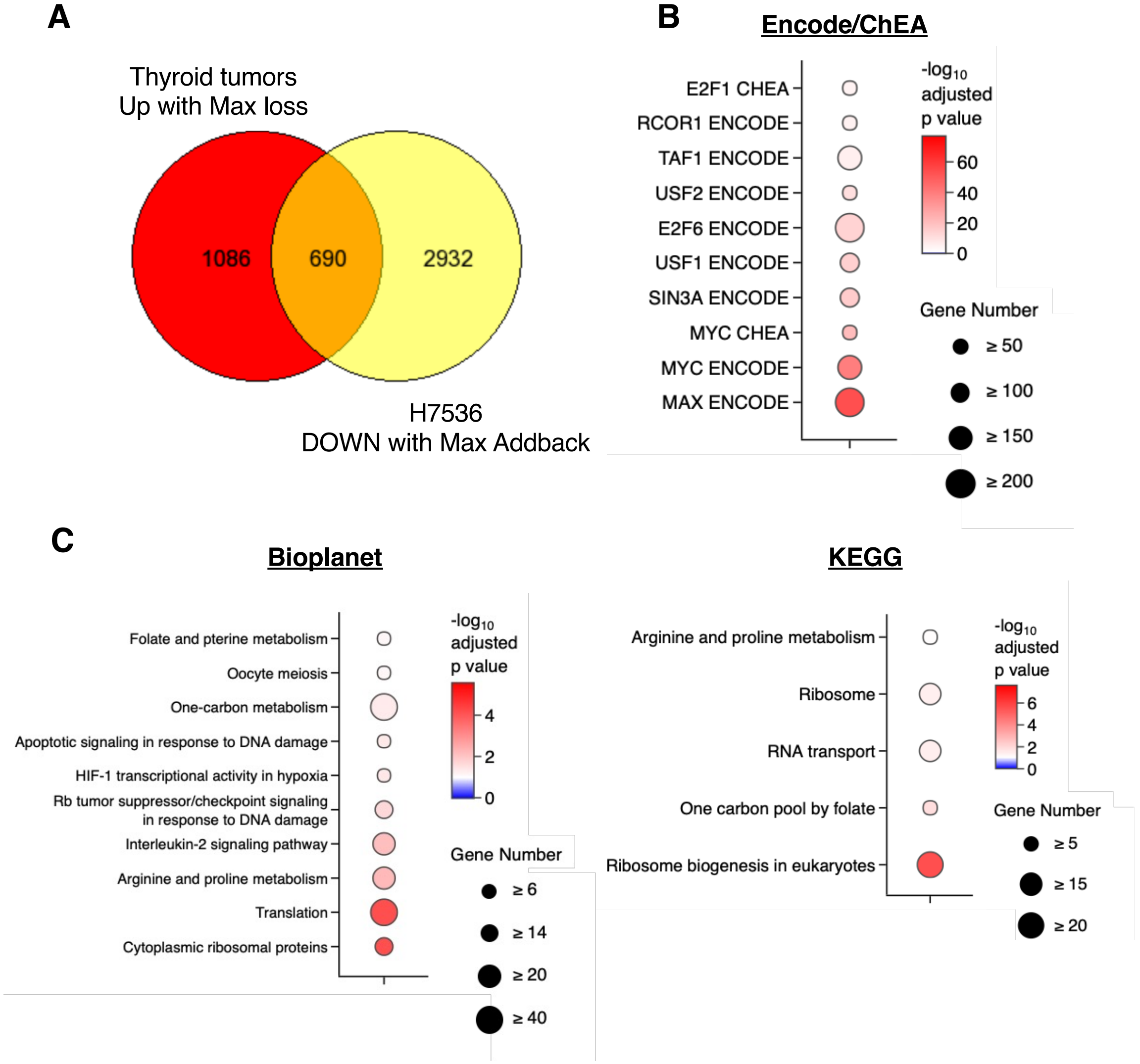


**Supplemental Figure 1. RNA-Seq data showing genes anti-correlated with MAX in thyroid cancer. A)** Venn Diagram showing genes with increased expression upon MAX loss (FDR <0.05) in EdgeR analyses comparing RPMax to RP thyroid tumors and genes with decreased expression upon doxycycline-induced return of MAX to H7536. **B, C, D)** Pathway enrichment analyses querying GO Biological Processes (F) and ENCODE/CHEA datasets (G) including the 50 core MAX-regulated genes from (Fig.3E).

**Fig.S2**

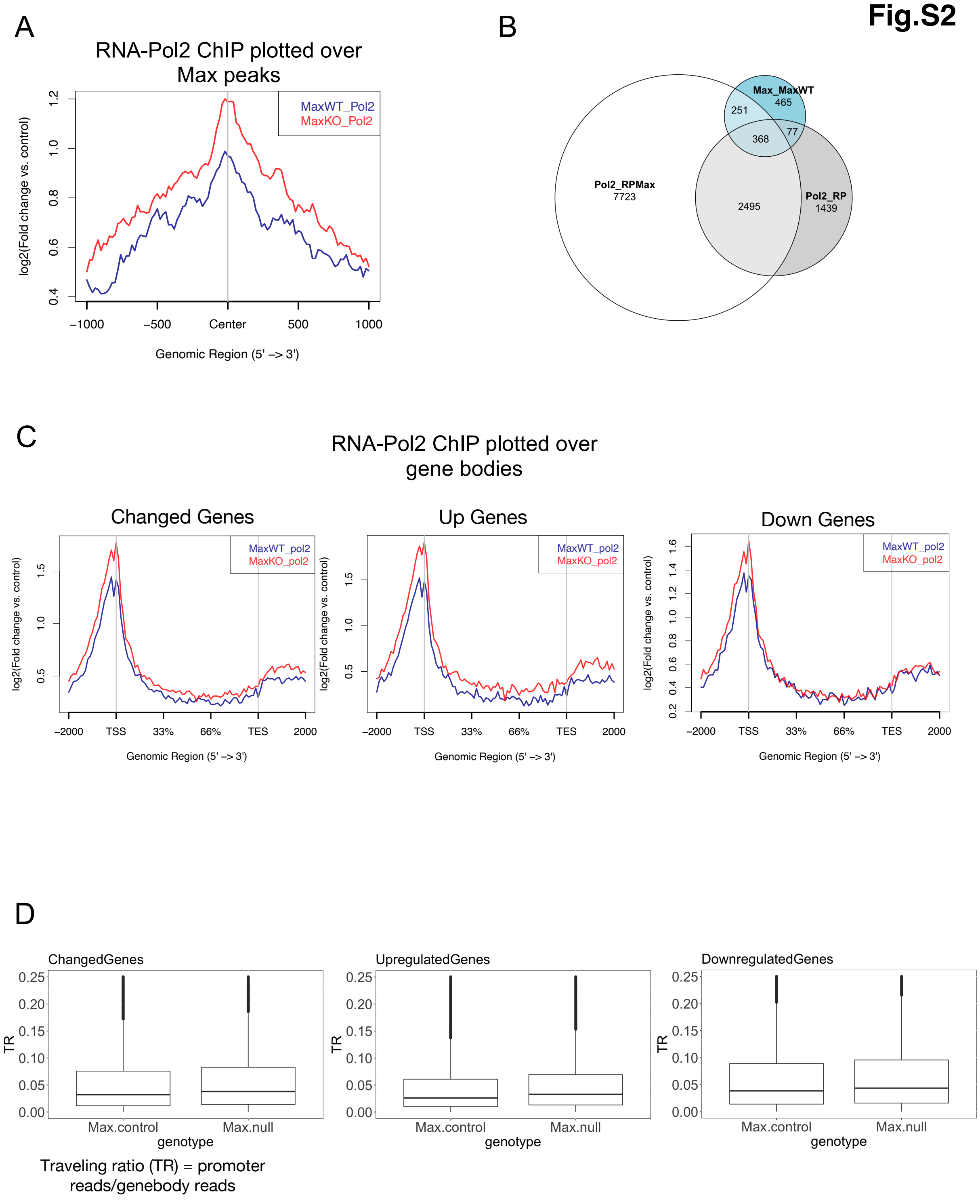
A
RNA-Pol2 ChIP plotted over Max peaks
MaxWT_Pol2
MaxKO_Pol2
log2(Fold change vs. control)
0.4
0.6
0.8
1.0
1.2
−1000
−500
Center
500
1000
Genomic Region (5' –> 3')
B
Max_MaxWT
465
251
368
77
Pol2_RPMax
7723
2495
Pol2_RP
1439
C
RNA-Pol2 ChIP plotted over gene bodies
Changed Genes
Up Genes
Down Genes
MaxWT_pol2
MaxKO_pol2
log2(Fold change vs. control)
0.5
1.0
1.5
0.2
0.4
0.6
0.8
1.0
1.2
1.4
1.6
−2000
TSS
33%
66%
TES
2000
Genomic Region (5' –> 3')
D
ChangedGenes
UpregulatedGenes
DownregulatedGenes
TR
0.00
0.05
0.10
0.15
0.20
0.25
Max.control
Max.null
genotype
Traveling ratio (TR) = promoter reads/genebody reads

**Supplemental Figure 2. RNA polymerase II genomic occupancy in RP and RPMax thyroid tumor cell lines. (A)** ChIP was performed using anti-RNA polymerase II (RNA-Pol2) antibody on 3 Max WT RP and 3 RPMax cell lines. After sequencing and alignment, aligned sequence files were merged by genotype (MaxWT or MaxKO). RNA-Pol2 is plotted over the center of MAX peaks (determined by peakcalls), with the coverage for MaxWT cells (blue), and MaxKO cells (red). **(B)** Venn diagram showing the overlap of RNA-Pol2 (Pol2_MaxKO and Pol2_MaxWT) and MAX peaks (Max_MaxWT). **(C)** Plots were generated for each genotype over all gene bodies of changed, up-regulated (Up Genes), and down-regulated (Down Genes) genes flanked by 2 kb of genomic region. **(D)** Traveling ratios were determined by dividing promoter mapped reads by gene body mapped reads for both MAX WT RP (Max.control) and MAX KO RPMax (Max.null) cells.

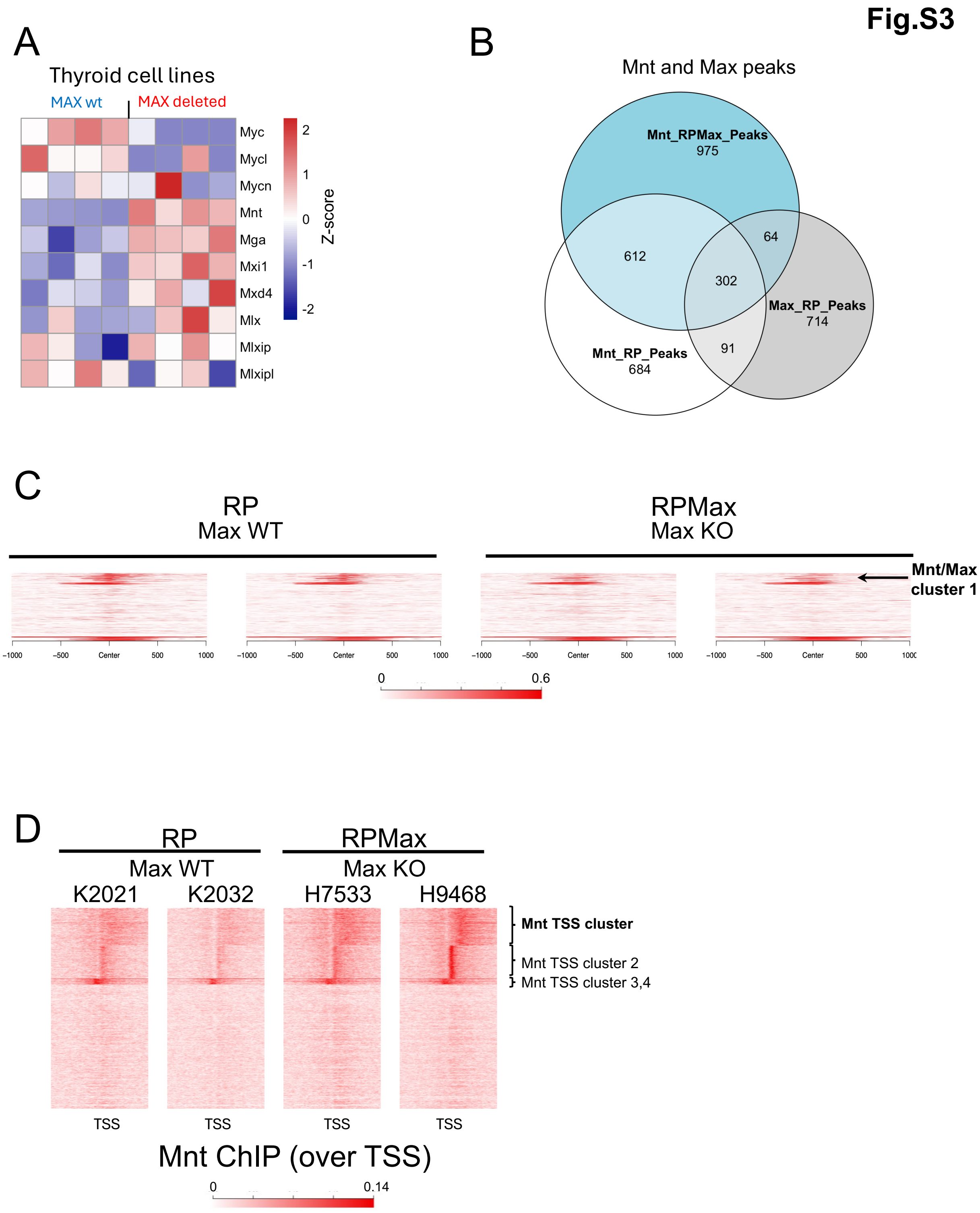

Fig.S3
A
Thyroid cell lines
MAX wt
MAX deleted
Myc
Mycl
Mycn
Mnt
Mga
Mxi1
Mxd4
Mlx
Mlxip
Mlxipl
2
1
0
-1
-2
Z-score
B
Mnt and Max peaks
Mnt_RPMax_Peaks
975
612
64
302
Max_RP_Peaks
714
91
Mnt_RP_Peaks
684
C
RP
Max WT
RPMax
Max KO
Mnt/Max cluster 1
-1000
-500
Center
500
1000
0
0.6
D
RP
Max WT
RPMax
Max KO
K2021
K2032
H7533
H9468
Mnt TSS cluster
Mnt TSS cluster 2
Mnt TSS cluster 3,4
TSS
Mnt ChIP (over TSS)
0
0.14

**Supplemental Figure 3. MYC network expression, MNT occupancy and peak clustering in RP and RPMax cells. (A)** Expression levels of mRNAs encoding Myc network genes determined by RNA-Seq in RP (MAX wt) and RPMax (MAX deleted) thyroid cell lines. The expression of each is normalized (by row) for each cell line. **B)** ChIP-seq was carried out using anti-MNT or IgG antibody on tumor-derived cell lines that are *Max* WT (RP) or *Max* KO (RPMax), and peak calls were performed. The Venn diagram illustrates the overlap between MNT peaks in RP cells (Mnt_RP_Peaks), RPM cells (Mnt_RPMax_peaks), and MAX peaks (MAX_RP_Peaks). **C)** K-means clustering of MNT occupancy over MAX peaks (determined by peak calls for MAX). Five clusters are identified, and one cluster (Mnt/Max cluster1) is shown in the heatmaps for two RP and two RPMax cell lines. **D)** K-means clustering of MNT occupancy over TSS of all genes. Five clusters are identified; one cluster (Mnt TSS cluster 1) is shown in the heatmaps for two RP and two RPMax cell lines.

**Fig.S4**

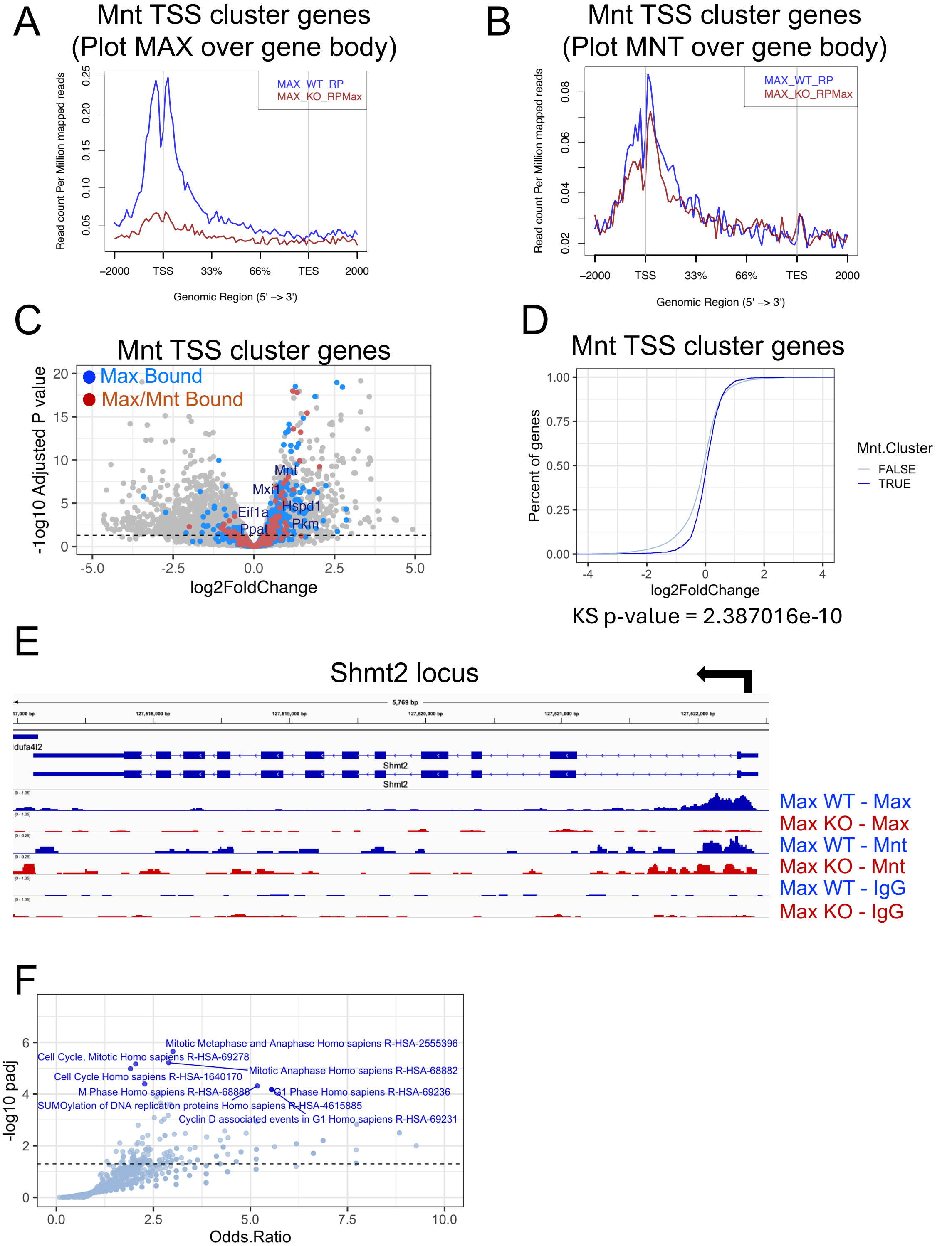
A
Mnt TSS cluster genes
(Plot MAX over gene body)
MAX_WT_RP
MAX_KO_RPMax
Read count Per Million mapped reads
0.05
0.10
0.15
0.20
0.25
−2000
TSS
33%
66%
TES
2000
Genomic Region (5' −> 3')
B
Mnt TSS cluster genes
(Plot MNT over gene body)
MAX_WT_RP
MAX_KO_RPMax
Read count Per Million mapped reads
0.02
0.04
0.06
0.08
−2000
TSS
33%
66%
TES
2000
Genomic Region (5' −> 3')
C
Mnt TSS cluster genes
Max Bound
Max/Mnt Bound
-log10 Adjusted P value
Mnt
Mxi1
Eif1a
Hspd1
Ppat
Pkm
-5.0
-2.5
0.0
2.5
5.0
log2FoldChange
D
Mnt TSS cluster genes
Percent of genes
Mnt.Cluster
FALSE
TRUE
log2FoldChange
KS p-value = 2.387016e-10
E
Shmt2 locus
5,769 bp
dufa4l2
Shmt2
Max WT - Max
Max KO - Max
Max WT - Mnt
Max KO - Mnt
Max WT - IgG
Max KO - IgG
F
Mitotic Metaphase and Anaphase Homo sapiens R-HSA-2555396
Cell Cycle, Mitotic Homo sapiens R-HSA-69278
Mitotic Anaphase Homo sapiens R-HSA-68882
Cell Cycle Homo sapiens R-HSA-1640170
M Phase Homo sapiens R-HSA-68886
G1 Phase Homo sapiens R-HSA-69236
SUMOylation of DNA replication proteins Homo sapiens R-HSA-4615885
Cyclin D associated events in G1 Homo sapiens R-HSA-69231
-log10 padj
Odds.Ratio

**Supplemental Figure 4. A subset of MAX/MNT bound genes are depressed with MAX loss**

**A-B)** Plots of genomic region over gene bodies (x-axis) vs normalized coverage (y-axis) of MNT TSS cluster 1 (see Figure S5D), which was determined to be differentially occupied by MNT. Coverage for RP cells (MAX WT, blue) is compared to that from RPM cells (MAX KO, red). Panel E shows MAX coverage, and Panel F shows MNT coverage. **C)** Volcano plot was generated plotting RNASeq gene expression changes comparing Max WT RP vs Max mutant RPM tumors (Log2FoldChange on the x-axis, -Log10 adjusted p value on y-axis). Genes with peaks that map to MAX bound sites (blue) and in MNT TSS cluster 1 (red) are shown. **D)** Cumulative distribution plots depicting the rank-order (y-axis, Percent of genes) vs log2FoldChange (x-axis) of RNASeq data (comparing Max KO vs Max WT tumors). MNT TSS cluster 1 genes (dark blue) and a similar sized set of genes determined to not be in an MNT occupied cluster (light blue) are shown. The difference between the distributions are statistically determined using Kolmogorov-Smirnov (KS) statistics. **E)** Genomic tracks showing differential occupation of the Mnt promoter in RP (Max WT) and RPMax (Max KO) cells as determined by ChIP-Seq. The antibodies against MAX, MLX and MNT used for ChIP are shown after the hyphen on the right. **F)** Enrichment analysis of genes found to be differentially occupied by Max and Mnt in Max WT tumor-derived thyroid cells (Mnt TSS cluster1 from S5D). Enrichment categories are plotted as the -log10 of the adjusted P value (y-axis, -log10 padj), and Odds.Ratio (x-axis).

**Fig.S5**

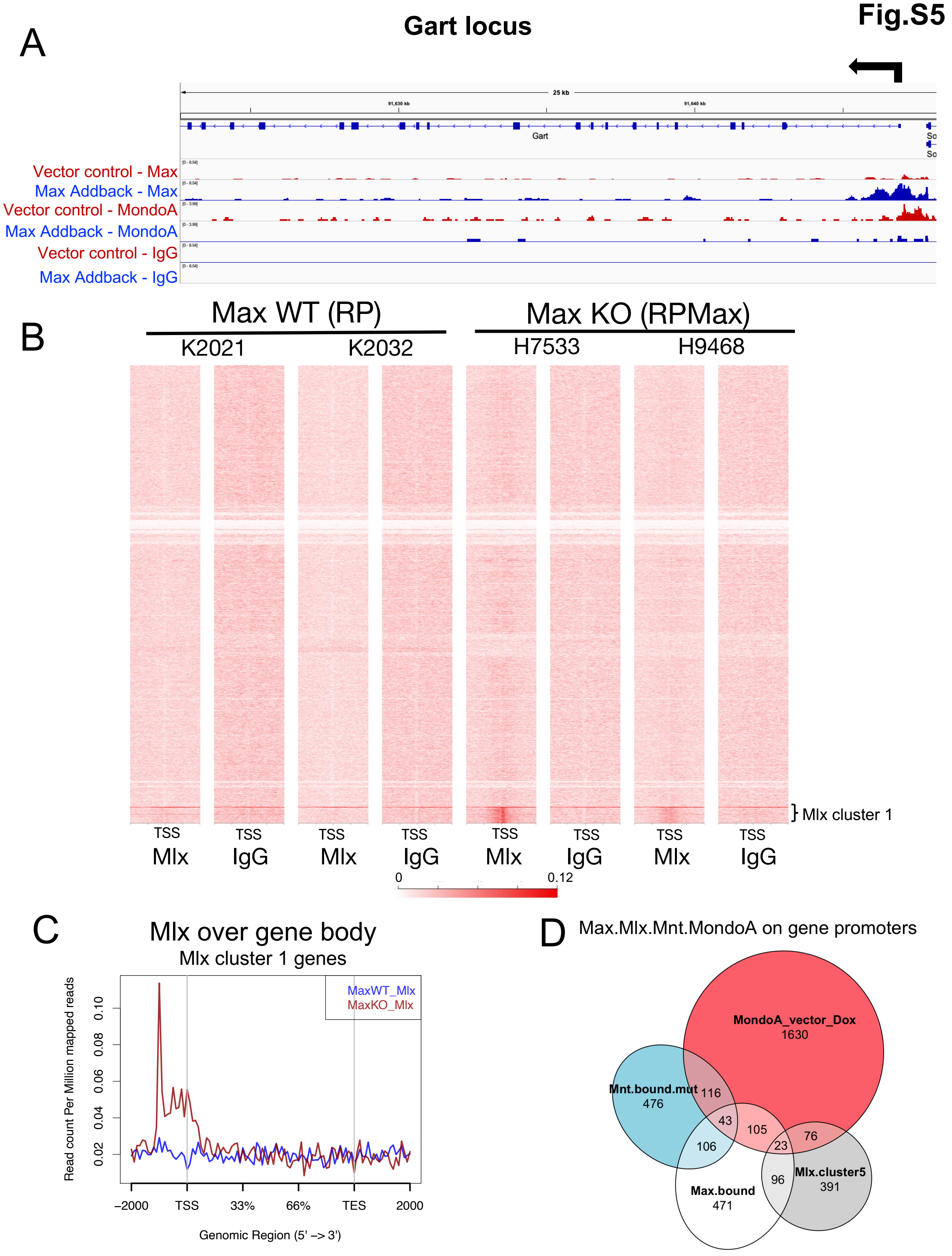
A
Gart locus
25 kb
Gart
Vector control - Max
Max Addback - Max
Vector control - MondoA
Max Addback - MondoA
Vector control - IgG
Max Addback - IgG
B
Max WT (RP)
Max KO (RPMax)
K2021
K2032
H7533
H9468
} Mlx cluster 1
TSS
Mlx
IgG
0
0.12
C
Mlx over gene body
Mlx cluster 1 genes
MaxWT_Mlx
MaxKO_Mlx
Read count Per Million mapped reads
0.02
0.04
0.06
0.08
0.10
–2000
TSS
33%
66%
TES
2000
Genomic Region (5' –> 3')
D
Max.Mlx.Mnt.MondoA on gene promoters
MondoA_vector_Dox
1630
Mnt.bound.mut
476
116
43
105
106
23
76
96
Max.bound
471
Mlx.cluster5
391

**Supplemental Figure 5. MondoA and Mlx occupancy at Max-regulated genomic loci. A)** Genomic tracks showing occupancy at the *Gart* locus by MAX and MondoA in *Max*null cells reconstituted with Max (Max-addback) or vector controls. The antibodies used for ChIP are shown after the hyphen on the left. **B)** Genomic occupancy by MLX in RP and RPMax tumor cell lines. ChIP was carried out using anti-MLX or IgG antibody on tumor-derived cell lines (2 RP and 2 RPMax). **C)** Plots of genomic region over gene bodies (x-axis) vs normalized coverage (y-axis) for the genes in Mlx cluster 1 (see Figure S5B). Coverage of Mlx for RP (MAX WT, blue) is compared to that from RPMax (MAX KO, red) cells. **D)** Venn diagram depicting the co-occupancy of gene loci by Max (Max_bound), Mnt (Mnt_bound_mut) and MondoA in *Max*-mutant cells (MondoA_vector_Dox), and Mlx in *Max*-mutant cells (Mlx_cluster_1).
